# AATF/Che-1, a new component of paraspeckles, controls R-loops formation and Interferon activation in Multiple Myeloma

**DOI:** 10.1101/2021.08.04.455054

**Authors:** Tiziana Bruno, Giacomo Corleone, Clelia Cortile, Francesca De Nicola, Valeria Catena, Francesca Fabretti, Svitlana Gumenyuk, Francesco Pisani, Andrea Mengarelli, Claudio Passananti, Maurizio Fanciulli

## Abstract

Multiple myeloma (MM) is a hematological neoplasm of plasma cells characterized by abnormal production of immunoglobulins. Che-1/AATF (Che-1) is an RNA binding protein involved in transcription regulation and is highly expressed in this malignancy. Here we experimentally show that Che-1 interacts with paraspeckle components, including the lncRNA NEAT1_2 (NEAT1), which serves as the seed for the maintenance of these structures. Che-1 and NEAT1 localize on R-loops, three-stranded RNA:DNA hybrids structures involved in DNA transcription and repair. Depletion of Che-1 produces a marked accumulation of RNA:DNA hybrids sustaining activation of a systemic inflammatory response. We provide evidence that high levels of Unfolded Protein Response (UPR) in MM cells induces RNA:DNA hybrids and an interferon (IFN) gene signature. We found that MM patients exhibit elevated R-loops levels and paraspeckle genes mRNAs increase linearly to MM progression. Strikingly, patients showing elevated IFN genes signature are associated with a marked poor prognosis. Overall, these findings delineate that elevated R-loops accumulation and inflammatory signaling may contribute to MM progression and that Che-1/NEAT1 plays an essential role in maintaining R-loops homeostasis by preventing excessive inflammatory signaling.

## Introduction

Multiple myeloma (MM) is a hematological malignancy characterized by the uncontrolled proliferation of transformed plasma cells in the bone marrow and the overproduction of immunoglobulins(1). This neoplasm is characterized by a high frequency of chromosomal aberrations, genomic instability (2–4), and intrinsic DNA damage (5). Despite continuing advances in the treatment of this disease, MM remains an incurable disease (1) and understanding the molecular mechanism underlying the progression and identification of key factors driving the aggressiveness is of fundamental importance.

Recent studies demonstrated that the protein Che-1/AATF (Che-1) promote MM transformation and progression by affecting chromatin accessibility and global transcription (6, 7). Che-1 is an RNA polymerase II-interacting protein highly conserved during evolution (8). During the last 20 years, it has been demonstrated that Che-1 is involved in many fundamental cellular processes, such as transcriptional regulation, cell-cycle and apoptosis control, cellular response to DNA damage and stress, and ultimately progression of many types of cancer (9–13).

Accumulating evidence indicates that transcription is regulated by the action of paraspeckles, nuclear bodies identified in the interchromatin space of the cells (14, 15). These structures are composed of specific RNA and at least 40 different proteins, essentially RNA binding proteins (RBP) expressed ubiquitously(15). The formation of the paraspeckle is mediated by the architectural long noncoding RNA (lncRNA) NEAT1_2 (NEAT1) which acts as a scaffold for the assembly of the components of this organelle (16). In addition to NEAT1, numerous RBPs such as NONO, SFPQ, FUS or RBM14 are essential for the formation and maintenance of paraspeckles (17). These organelles dynamically control the shuttling of specific proteins and mRNAs between this compartment and the nucleoplasm. Nevertheless, NEAT1 interacts with chromatin and localizes at transcription start sites (TSS) of active genes, leading to hypothesize a direct involvement of this lncRNA on transcription (18).

During transcription, the nascent RNA binds with the template strand of DNA, removing the complementary DNA strand, thus leading to the formation of RNA:DNA hybrids. These three-stranded nucleic acid structures, composed of RNA:DNA hybrids and the associated non-template single-stranded DNA are called R-loops (19, 20). Failure to control the generation of these structures can lead to transcription blockage and the generation of DNA damage and genomic instability. Numerous mechanisms are involved in the correct regulation of R-loops, and their accumulation appears to be related to numerous neurodegenerative and oncological pathologies (19–22).

In this study we show that Che-1 interacts with numerous paraspeckle components, in particular with lncRNA NEAT1. Che-1 and NEAT1 colocalize on the DNA, and NEAT1 modulates the presence of Che-1 on chromatin, regulating the resolution of R-loops. Depletion of Che-1 or NEAT1 produces an accumulation of these structures and activation of inflammatory pathway. Notably, we demonstrate that hybrid structures increase in MM patient cells and observed a linear relationship between the progression of MM and the expression of paraspeckle and IFN response genes. Finally, our findings suggest a marked worse prognosis in MM patients showing elevated activation of IFN response.

## Material and Methods

### Cell lines and cell culture treatment

Human multiple myeloma cell lines Kms27, U266, and RPMI-8226 were cultured in Optimem (Thermo Fisher Scientific) supplemented with 15% inactivated Fetal Bovine Serum (FBS) (Thermo Fisher Scientific), 2mM glutamine and 40μg/ml gentamicin. All cell lines were cultured at 37°C, in a humidified atmosphere with 5% CO2 and mycoplasma contamination was periodically checked by polymerase chain reaction (PCR) analysis. All the cell lines were not listed in the ICLAC database of commonly misidentified cell lines and were authenticated at The San Martino Hospital Genomics Facility using short tandem repeat DNA profiling. The presence of the Epstein-Barr virus (EBV) was detected by PCR analysis. ISRIB (Sigma SML0843) and Thapsigargin (Sigma T9033) were resuspended in DMSO as instruction and added to cell medium at concentration and for a time in according to the effect evinced by a dose curve.

### Transfection with siRNA and DNA constructs

All the transfection experiments were carried out using the Lipofectamine 3000 reagents according to the manufacturer’s instructions (Thermo Fisher Scientific). Cells were analyzed 48 hours after transfection, and the efficiency of transfection was determined by RT-qPCR, immunoblotting or immunofluorescence. Stealth siRNA oligonucleotides targeting Che-1 (siChe-1, cat. n. 1299003– HSS120158 and siChe-1 b, cat. n. 1299003–HSS120159) or a control sequence (siControl, cat. n. 12935300) were purchased from Thermo Fisher Scientific. pEGF-RNaseH1 expression vector was purchased from Addgene (plasmid #108699). RelA expressing vector and NFkB-luc reporter were a gift Dr. Karin from University of San Diego, La Jolla (CA). LNA oligonucleotides were provided by Exiqon (Vedbaek, Denmark). Custom Long Non-Coding LNA GapmeRs were custom-designed and purified by HPLC followed by Na+-salt exchange and lyophilization. LNA GapmeRs were used at 100nM.

### Human specimens

MM patient samples were collected as part of routine clinical examination. The study was approved by the Regina Elena Cancer Institute Ethics Committee (CE 422/14) and written informed consent to participate in this study was provided by all subjects. Bone marrow aspirates from healthy donor and pathological patients were enriched for plasma cells by magnetic cell separation using a human CD138 positive selection and Macs Separator kits (Miltenyi Biotec, Germany).

### Western blot and coimmunoprecipitation

Immunoblotting and coimmunoprecipitation were performed as previously described(23). Detailed information for all antibodies is provided in Supplemental Table S2.

### Luciferase assay

Cell extracts were prepared and assayed for luciferase activity according to the manufacturer’s instructions (Promega, WI) using the GloMax Luminometer (Promega, WI). Total protein quantification in the extracts was determined by Bradford assay, and luciferase activity of equal amounts of proteins was determined.

### RNA isolation and quantitative real-time PCR

Total RNA was extracted from the same number of viable cells using QIAzol Lysis Reagent (Qiagen). cDNA was synthesised from equal amount of RNA by reverse transcription using M-MLV reverse transcriptase (Thermo Fisher Scientific) and a mixture of random primers (Thermo Fisher Scientific). This single-stranded cDNA was then used to perform quantitative real-time PCR (RT-qPCR) with specific primers using a PowerUp SYBR Green Master Mix (ThermoFisher) on a 7500 Fast Real-Time PCR System (Applied Biosystems) following the manufactures’ instructions. The Ct was measured during the exponential amplification phase, and the amplification plots were analyzed using the software v2.0.6 (Applied Biosystems). Relative fold changes were determined by comparative threshold (ΔΔCT) method using *β-actin/RPL19* genes as endogenous normalization control (24). Data are presented as mean ± SD of three independent experiments, performed in triplicate. Specific primers employed in RT-qPCR amplifications are listed in Supplemental Table S3.

### Proximity ligation assay (PLA)

Cells were centrifuged mounted on glass slides using a Thermo Shandon Cytospin 2 (Thermo Fisher Scientific) and fixed in 4% formaldehyde for 10 min and permeabilized with 0.1% Triton X- 100 in PBS for 5 min at room temperature. In *situ* proximity ligation assay (PLA) was performed using the Duolink InSitu Reagents (Sigma-Aldrich) according to the manufacturer’s instructions. Images were acquired using a fluorescence microscope with a 40X objective (Zeiss, Germany) and processed with AxioVision Software.

### Mass-spectrometry (MS)

Mass-spectrometry (MS) analysis was performed immunoprecipitating Che-1/AATF from total extracts of Kms27 cells. Immunoprecipitation with normal IgGs was performed as negative control. Co-immunoprecipitated complexes were eluted by incubation in 5% SDS in PBS at 95°C and stored at −80°C prior LC-MS analysis. MS processing was performed as previously described (25)

### RNA immunoprecipitation (RIP) with crosslinking

Cells were crosslinked with formaldehyde (F.A.) solution (50 mM HEPES-KOH pH 7.5, 100 mM NaCl, 1 mM EDTA, 0.5 mM EGTA, 11% formaldehyde, ddH2O) 1% final concentration, 10 min, RT. Nuclei were isolated and resuspended in lysis buffer (Tris–HCl pH 7.5 50 mM, EDTA 1 mM, SDS 0.5%, DTT 1 mM), using 200 μl for each IP sample, and sonicated to obtain a smear not higher than 500 bp. Lysate was treated with DNase (DNAfree, Ambion) and diluted with 400 μl of correction buffer (NP-40, 0.625%, DOC, 0.312%, MgCl2, 5.6 mM, Tris–HCl pH 7.5, 47.5 mM, NaCl, 187.5 mM, glycerol, 12.5%, DTT 1 mM). IP was carried out overnight at +4°C with Che-1/AATF (Bethyl) or S9.6 (Kerafast) antibodies, while normal mouse IgG (Bethyl) functioned as a negative control. IP washing and proteinase K digestion were carried out as above, crosslinking was reversed by incubation at 70°C for 30 min, and RNA was recovered by TRIzol extraction. Immunoprecipitated RNA was tested by RT-qPCR analysis to verify the presence of lncNEAT1 using specific primers (Supplemental Table 3) and were calculated by standard curve method and fold of enrichment over IgG.

### FISH

Cy5 labeled probes for human NEAT1 (Stellaris RNA Biosearch Techonologies) were used for FISH experiments. For co-localization experiments we follow the protocol for IF+FISH in Suspension Cells with minor modification. After the permeabilization, we blocked the cytocentrifuge with BSA 5% in PBS1x for 1 hour, followed by a wash in PBS 1x for ten minutes, then we proceeded in according with manufacture’s protocol. The images were captured and analyzed by Zeiss LSM 880 with Airy scan confocal laser scanning microscope by 63X/1.23 NA oil immersion objectives. Lasers 670 nm were used to excite the fluorophores. The Zeiss Zen control software (Zeiss, Germany) was used for image analysis.

### Immunofluorescence

Cytocentrifuged cells were fixed in 4% formaldehyde for 10 min and then permeabilized with 0.1% Triton X-100 in phosphate-buffered saline (PBS) for 5 min at room temperature. Cells were subsequently stained for 2 h with primary antibodies, after which they were rinsed three-times with PBS and stained for 45 min with Alexa-Fluor-594- and Alexa-Fluor-488-conjugated anti-rabbit or anti-mouse secondary antibodies (Thermo Fisher Scientific). Nuclei were visualized by staining with 1 μg/ml Hoechst dye 33258 (Sigma-Aldrich).

For DNA-RNA hybrids staining, cells were cytocentrifuge and then fixed in ice-cold methanol for 10 minutes at -20°C. The slides were then quickly washed in PBS 1X and incubated for 1h in PBS 1x 3% BSA to prevent unspecific interaction. Primary antibodies S9.6 was incubated for two hours at room temperature (RT) in the dark. After three washes with PBS 1x the slides were incubated at RT with secondary anti mouse antibodies. Slides were then washed with PBS 1X and mounted using Duo link in situ mounting medium with DAPI (Sigma). Quantification of the hybrids signal intensity per cell were performed by counting 100 cells per condition in each experiment and analysis were made using ImageJ software.

### RNase H treatments

For IF experiments, cells were cytocentrifuged and permeabilized with PBS 1x plus 0,5% Tween 20 for 5 minutes at RT. Cells were then treated with 10U of RNase H (NEB), in PBS 1x for 1 hour at RT. For Dot Blot and DRIP seq experiments RNase H treatments were performed on genomic DNA with 10U of the enzyme for 1h at 37°C.

### Dot Blot

Genomic DNA was extracted according to DRIP protocol. Serial dilutions of DNA were double spotted on a nitrocellulose membrane and crosslinked with UV light (700 mJ/cm2). One part of the membrane was blocked with TBS-Tween 0,1% and 5% non-fat dry milk (NFDM) for 1 hour and then incubated with S9.6 antibody diluted to 0.5 μg/ml in TBS-Tween plus 2% NFDM. After washing, membrane was incubated with anti-mouse secondary antibodies, further washed, and developed with ECL techniques. The other part of membrane was incubated with Methylene Blue (Sigma M4159 0,2% (w/v) in 0,4M sodium acetate:0,4M acetic acid) for 15 minutes, as loading control, and at the end of the incubation washed with water. Both the membranes were acquired and quantified using Alliance Mini HD6 system by UVITEC Ltd, Cambridge, equipped with UVI1D Software (UVITEC, 14–630275).

### ChIP-seq

ChIP-seq experiments were performed following ChIPmentation protocol (7) in duplicate for Che-1/AATF. The final libraries were controlled on an Agilent 2100 Bioanalyzer (Agilent Technologies) and sequenced on a NextSeq 500 (Illumina, CA) using 50 cycles single-end mode.

### RNA-Seq

Total RNA was extracted from the same number of viable cells using QIAzol Lysis Reagent (Qiagen). ExFold RNA Spike-In Mixes (ERCC, Thermo Fisher Scientific) were added to each sample for normalizing gene expression. RNA libraries for sequencing were generated in triplicate by using the same amount of RNA for each sample according to the Illumina TruSeq Stranded Total RNA kit with an initial ribosomal depletion step using Ribo Zero Gold (Illumina, CA). The libraries were quantified by RT-qPCR and sequenced in paired-end mode (2x75 bp) with NextSeq 500 (Illumina, CA).

### ChIRP-seq

ChIRP was performed as described (26). 20 million cells were used for each condition. The oligonucleotides used for NEAT1 immunoprecipitation are listed in Supplemental Table S3. Biotinylated oligonucleotides were recovered using Dynabeads® MyOne™ Streptavidin C1 (Invitrogen). After the washing steps, all the recovered material was used for DNA purification and subsequently utilized for the construction of the library by Accel-NGS 2S Plus DNA library kits (Swift biosciences) in according to the manufacturer’s instructions.

### DRIP-seq

DRIP-seq assays were performed as previously described (27). Briefly, nucleic acids were extracted from Kms27 cells by SDS/proteinase K treatment at 37°C overnight followed by phenol– chloroform isoamyl extraction and ethanol precipitation at room temperature. The harvested nucleic acids were digested for 24 h at 37°C using a restriction enzyme cocktail (50 units/100 μg nucleic acids, each of BsrGI, EcoRI, HindIII, SspI, and XbaI) in the New England Biolabs CutSmart buffer with 2 mM Spermidine and 1× BSA. Digested DNAs were cleaned up by phenol–chloroform isoamyl extraction followed by treatment or not with RNase H (10 units/100 μg nucleic acids) overnight at 37°C in the New England Biolabs RNase H buffer. DNA/RNA hybrids from 4.0 μg digested nucleic acids, treated or not with RNase H, were immunoprecipitated using 5 μg of S9.6 antibody (Kerafast) and 20 μl of Protein G Dynabeads (Thermo Fisher) at 4°C overnight in IP buffer (10 mM NaPO4, 140 mM NaCl, and 0.05% Triton X-100). The beads were then washed four times with IP buffer for 10 min at room temperature, and the nucleic acids were eluted with elution buffer (50 mM Tris–HCl, pH 8.0, 10 mM EDTA, 0.5% SDS, and 70 μg of protease K) at 55°C for 1 h. Immunoprecipitated DNA was then cleaned up by a phenol–chloroform extraction followed by ethanol precipitation at - 20°C for 1h. The quantity of the immunoprecipitated material was determined by Qubit 2.0 fluorometer (Life Technologies). About 10 ng of the immunoprecipitated DNA was used to prepare the libraries for sequencing by following the manufacturer’s instructions including DNA and repairing, adaptor ligation, and amplification (Illumina, CA). The libraries were then sequenced using the NextSeq 500 system (Illumina, CA).

## Computational Methods

### ChIRP-seq and ChIP-seq data processing

Reads of each experiment were quality controlled with FASTQC v0.11.9 (http://www.bioinformatics.babraham.ac.uk/projects/fastqc/) combined with MultiQC v.1.9(28) and aligned to the reference genome hg19 using bowtie2 (29) v.2.3.5.1 setting –local mode with default parameters. The generated sequence alignments were stored into SAM format files and converted into BAM format, sorted, and indexed with SAMtools(30) v.1.2. The duplicated reads were removed by utilizing MarkDuplicated tool available in GATK4 suite v.4.1.8.1. Peaks were called using the function callpeak of MACS2 v.2.2.6 tool (parameters: --format AUTO --broad -B -q 0.1 -g hs), and those matching backlisted regions of hg19 were removed. Peaks stored in bedGraph format files were sorted and converted in bigWig format with bedGraphToBigWig(31) v.4. Significant signals were annotated by using the function annotatePeaks.pl of HOMER (32) v.4.11 tool.

### Heatmap and enrichment profiles of ChIRP-seq, ChIP-seq, DRIP-seq profiles

Heatmap profiles were obtained by respectively using the function included in DeepTools v.3.5.0 computeMatrix (parameters: scale-regions -a 5000 -b 5000 –region Body Length 5000 -- skipZeros –missingDataAsZero) and plotHeatmap (parameters: --sortRegions descend --sortUsing max --colorMap OrRd --zMin 0 --zMax 15 --dpi 300). Visualization of peak profiles were obtained using bigwig files of each experiment provided as input to Integrative Genomics Viewer (IGV).

### NEAT1 and Che-1 co-localization identification

The identification of the co-localizing sites between Che-1 and NEAT1 relies on a multistep process which include the generation of a master list using TSSs coordinates reported in the Ensembl release 103 (33) and referred to the GRCh37 reference genome. The master list was obtained using the biomart tool available on the Ensembl website and refined by removing mitochondrial coordinates and patches. A total of 92.226 TSSs were retained in the final master list. Each set of coordinates was extended by 2kb, both upstream and downstream. The master list was populated at each TSS site with Che-1, and NEAT1 read enrichment computed by using the multicov function of bedtools v2.29.2 tool and using ChIRP-seq and ChIP-seq deduplicated bam files as input. The obtained reads counts were quantile normalized after log2-counts per million transformation with the function voom of LIMMA (34) v3.44.3 package in R.

The identification of Che-1 and NEAT1 co-localized signal was obtained by using read enrichment cut-offs independently as follows: 1) The normalized reads count assigned to each TSS of both experiments were used to determine two independent series of cut-offs by dividing each distribution of normalized read counts in 15 quantile sections. 2) Iteratively, our algorithm scans each peak from the 1st (lowest enriched) quantile to the 15th quantile (top enriched) in incremental order. Each iteration includes the peaks being part of the previous quantiles. At each iteration assigns a value of 1 to the peak included in the quantile group, and a value of 0 in the case is not called. A signal is considered co-localized when in both experiments, the assigned value is 1. Each iteration is concluded with the calculation of a Jaccard coefficient score. 3) The number of co-localizing sites and Jaccard score obtained from each iteration were independently plotted with an in-house R script. The intersection of the two lines was considered the point of threshold for co-occurrence selection. The point of intersection corresponded to the 11th iteration.

### RBPs binding to NEAT-1-protein analysis

*BednarrowPeak* and *bigWig* files of 120 RBPs eCLIP profiles generated from K562 cell line and aligned to HG19 reference genome were downloaded from Encode Project database (v113). Peaks intersecting the NEAT1 domains (left domain: chr11:65190234-65198234; central domain: chr11:65198234-65206834; right domain: chr11:65206834-652129) (PMID: 29932899) with a length greater than 500bps were retained for further analysis. The classification of peaks was plotted by using ggplot2^8^ R package.

### DRIP-seq data processing

DRIP-seq data were quality checked by using FASTQC v0.11.91, forwarded to bowtie23 v.2.3.5.1(parameters: -p 6 -t --local) and aligned to the reference genome hg19. The alignments were saved in SAM format files and converted into BAM format which were subsequently sorted and indexed by using SAMtools4 v.1.2. Peaks were called with the function callpeak of MACS2 v.2.2.6 tools (parameters: --broad --format AUTO -B -q 0.01) and after sorting, were converted from bedGraph to BigWig format by using bedGraphToBigWig (31) v.4 tool. RNAse H treated sample bam file was used as input in the peak calling process.

### RNA-seq analysis

NEAT1 q vs. GapmerNEAT-1 RNA-seq and Che1 vs. siChe-1 RNA-seq were independently analysed by using the Kallisto v.0.46.0(35) and Sleuth v.0.30.0 (36) pipeline. Sequencing quality of each sample was assessed with FASTQC v0.11.91 and summarized with MultiQC v.1.92. Read quantification was performed with the function quant of Kallisto tool (parameters: -b 100). The pseudoalignment was executed against the reference genome GRCh37 downloaded from NCBI and subsequently indexed with the function index available in Kallisto.

The Kallisto results were processed and normalized through the function sleuth_prep with gene-level resolution (parameters: extra_bootstrap_summary=T, gene_mode = TRUE). The gene-transcripts correspondence map was retrieved through the functions useMart and getBM of biomaRT R package(37). Those correspondences were annotated based on grch37 of Ensembl release 103(33). Differential analysis was conducted by fitting full and reduced models through the function sleuth_fit with the formula parameter respectively set to full and reduced. The processed reads counts were converted in transcript per million measurements through the function sleuth_to_matrix of Sleuth (36) R package.

Principal component analysis (PCA) was executed pre- and post-normalization. Pre-normalized PCA was executed using the function prcomp of stats v.4.0.2 R package (http://www.r-project.org/index.html) and plotted through the function autoplot of ggfortify v.0.4.11 R package. Post-normalized PCA was carried out with the function plot_pca of Sleuth (36) package in R.

Wald test was performed with the function sleuth_wt of Sleuth (36) R package. The beta-value and q-value obtained from the test were used to discriminate between genes significantly up (b>0.7 and qval<0.05) or down (b<-0.7 and qval<0.05) regulated.

Functional enrichment analysis of differential expressed genes (DEGs) was executed through GSEA online portal released in July 2020 (http://www.gsea-msigdb.org) by interrogating the MSigDB Hallmarks collection. Pathway’s perturbation analysis was performed by running PROGENy (38) v.1.10.0 R package providing RNA-seq data as input.

### Transcriptome Analysis of the CoMMpass dataset

Raw gene counts of CoMMpass dataset (IA15 release) were downloaded from the MMREF Research Gateway Portal, table name MMRF_CoMMpass_IA14a_E74GTF_Salmon_Gene_Counts. We selected a cohort of 687 samples having assigned the relative ISS score and the transcriptome. Gene counts of selected samples were normalized through a multistep workflow using edgeR(39) functions as follows DGEList, estimateCommonDisp, estimateTaqwiseDisp, calcNormFactors with TMM method. The normalized matrix was populated with the cpm value to each gene within every patient. To identify genes differentially transcribed between ISS scores we iteratively screened each gene and then computing the Mann-Whitney test for each ISS combination providing the relative cpm value for each patient. Only genes differentially enriched (p-value<0.05) in at least one ISS combination were retained in the final matrix for further use.

### Enrichment of paraspeckles genes in disease progression

Paraspeckles genes (15) (*NEAT1, AATF, NONO, HNRNPK, DAZAP1, HNRNPH3, SRSF1, SFPQ, FUS, RBM14, TAF15*) were selected from the normalized matrix and plotted by the relative ISS score associated to each patient through ggpubr R package.

### IFN gene signature analysis

We investigated the expression of genes associated to the interferon gamma (IFNG) and interferon alpha (IFNA) pathway. We generated a customized list of 198 IFN genes which were collected from *Reactome, Signal Transduction Knowledge Environment, Biocarta, WikiPathways* stored in GSEA website (https://www.gsea-msigdb.org/gsea/msigdb/search.jsp) at the date of January 2021. Only IFN genes significantly up-regulated between the cohort of patients at ISS-1 and ISS-3 were retained for further analysis. A gene was considered significantly up regulated if the p-value obtained from the Mann-Whitney test was lower than 0.05 and the median cpm value of the whole cohort at ISS-3 was higher than ISS-1. The final selected IFN signature was composed by 29 genes which were consistently upregulated at ISS-3 compared ISS-1 and were used in the further analyses.

### IFN score assignment

An IFN score was assigned to each patient in the CoMMpass cohort. IFN score ranges from 1 to 3 respectively 1=Low,2=Medium, 3=High IFN genes expression. To define the IFN score we 1) calculated the value of the third quartile of each normalized gene count distribution within the CoMMpass cohort (N= 687) as independent cut-off for defining a gene as highly expressed. A patient was considered a high expressor of a gene when the relative gene expression was beyond the third quartile enrichment. Patients exhibiting less than 5 gene highly expressed were included in the IFN low group, those exhibiting between 5 and 13 genes were assigned to the IFN medium and those with more than 13 genes assigned to IFN high group. The frequency of patients in each group was plotted by the relative ISS with an in-house R script.

### Survival analysis

Two different survival analyses were conducted to reveal the role of IFN pathway in the MM disease progression. The overall survival (OS) information for the analysis was retrieved from table named ‘MMRF_CoMMpass_IA15_STAND_ALONE_SURVIVAL.csv’ stored on MMREF Research Gateway Portal. Patients with OS greater than 65 months were excluded (n=56) from the analysis. Survival curves were estimated with the KaplaneMeier product-limit method and compared by log-rank test. Univariate Cox regression analyses were carried out to identify potential predictors of survival.

The first survival curve examines the group of patients exhibiting high IFN score (n=71) versus the patients assigned to low and medium IFN score groups(n=614), defined as described above.

The second survival curve examine two groups of patients of similar size (IFN-low:341 vs IFN-high: 291). Patients were ranked based on the number of highly expressed gene involved in IFN pathway. Then, the median value of the ranking was set as threshold to divide the population in high vs low IFN signature.

Both survival analyses were carried out by using survfit and Surv functions in survival v.3.1.12 R package. The Kaplan-Meier curves were generated by utilizing the function ggsurvplot of survminer v.0.4.819 R package and both of them had significance equal to log-rank p-value<0.001.

### Multivariate analysis

The patients used for the survival analysis were classified according to: type of therapy (V-based, K-based, combo K/IMIDs-based, combo V/IMIDs/K-based, combo IMIDs/K-based, IMIDs- based), autologous stem-cell transplantation (ASCT; yes/no), presence or absence of translocations and copy-number abnormalities (CNAs). The definition of variables is accurately described in Supplementary Table 3. The translocations taken into consideration were those occurring at IgH and IgL locus (40) [t(11;14), t(4;14), t(14;16) t(4;20)] specifically found in MM disease, and were evaluated using calls on WGS long-insert data downloaded from MMREF Research Gateway Portal, table name MMRF_CoMMpass_IA15a_LongInsert_Canonical_Ig_Translocations. Nonsynonymous alterations occurring in a customized set of 21 genes (41) knowing to be mutated in MM were evaluated utilizing WGS long-insert data downloaded from MMREF Research Gateway Portal, table name MMRF_CoMMpass_IA15a_All_Canonical_NS_Variants_ENSG_Mutation_Counts.

Univariate Cox analysis of time to death was performed for each variable in the Table 2 by using the functions coxph and Surv of survival v.3.1.12 R package. Those variables resulted significant (pvalue≤0.06) at univariate analysis (IFN, ASCT, ISS, Therapy, KRAS, TP53, LTB, FGFR3) and two translocations essential in MM (MYC-translocation, CCND1-translocation) were used to carry out the multivariate Cox analysis of time to death. The statistical results were reported in Table 1. We fitted multivariate analysis by utilizing the functions coxph and Surv of survival v.3.1.12 R package with parameter na.action set to na.exclude. The multivariate analysis results were visualized with the function ggforest of survminer v.0.4.8 R package.

**Table 1:**
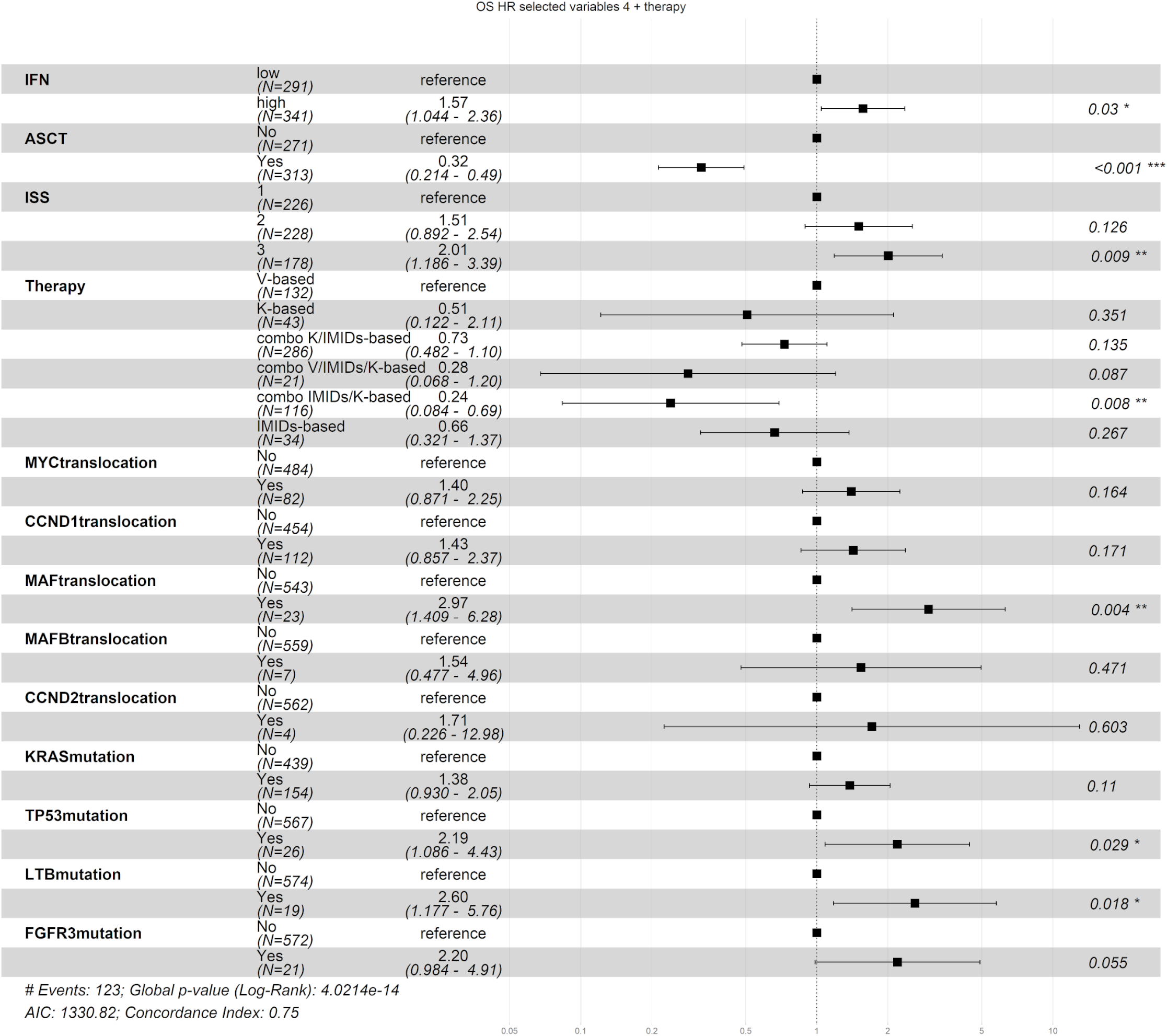
Multivariate analysis of Interferon upregulated patient cohort. Table including forest plot illustrating the multivariate Cox regression analysis for OS in the MM compass cohort with IFN signature selection expression on a continuous scale. The multivariate Cox model included 632 patients. 13 variables were considered to perform the analysis.

## Results

### Che-1 interacts with paraspeckle components

Che-1 is a nuclear protein involved in transcription regulation and cell proliferation of MM. To shed more light on its role in this disease, we performed co-immunoprecipitation experiments coupled with a mass-spectrometry analysis (Figure 1A). Mass spectrometry revealed that ∼600 proteins were significantly enriched in Che-1 precipitates. As expected from its role in ribosomal RNA maturation, the Che-1 interactome contains numerous proteins involved in pre-rRNA processing (42–44) and several ribosomal proteins (42). Notably, among other proteins interacting with Che-1, we identified many essential components of the paraspeckle (15) (Figure 1B), and these results were further confirmed by a previous mass-spectrometry analysis performed in HeLa cells (42) (Supplementary Figure S1A), thus suggesting a conserved role of Che-1/paraspeckles protein interaction in multiple model systems. To validate these results, we evaluated the binding of Che-1 to two important components of the paraspeckle, NONO and SFPQ. Reciprocal endogenous co-immunoprecipitations showed a strong interaction of Che-1 with both these proteins (Figure 1C) and proximity ligation assays (PLA) confirmed these results, showing discrete spots by fluorescence microscopy in Kms27 and RPMI8226 MM cells (Figure 1D). To investigate a direct binding of Che- 1 with NEAT1 we integrated in our framework data from the largest database of RBP binding to date (45). We leveraged our ability to infer RBPs/NEAT1 binding to date in immortalized leukemia and liver hepatocellular carcinoma cell lines (K562, HepG2) which confirmed the binding of Che-1 occur to NEAT1 5’ and 3’ functional domains (Figure 1E and Supplementary Figure S1B). In line with the model proposed by Yamazaki et al. (46), 3 fundamental components of the paraspeckle core, NONO, SFPQ and FUS contribute to paraspeckle assembly and stabilization by binding to both the middle and peripheral domains overlapping with Che-1 signal. These results were confirmed in two MM cell lines by RNA immunoprecipitation (RIP) and co-localization assays, clearly demonstrating a direct interaction between NEAT1 and Che-1 (Figures 1F and 1G).

**Figure 1:**
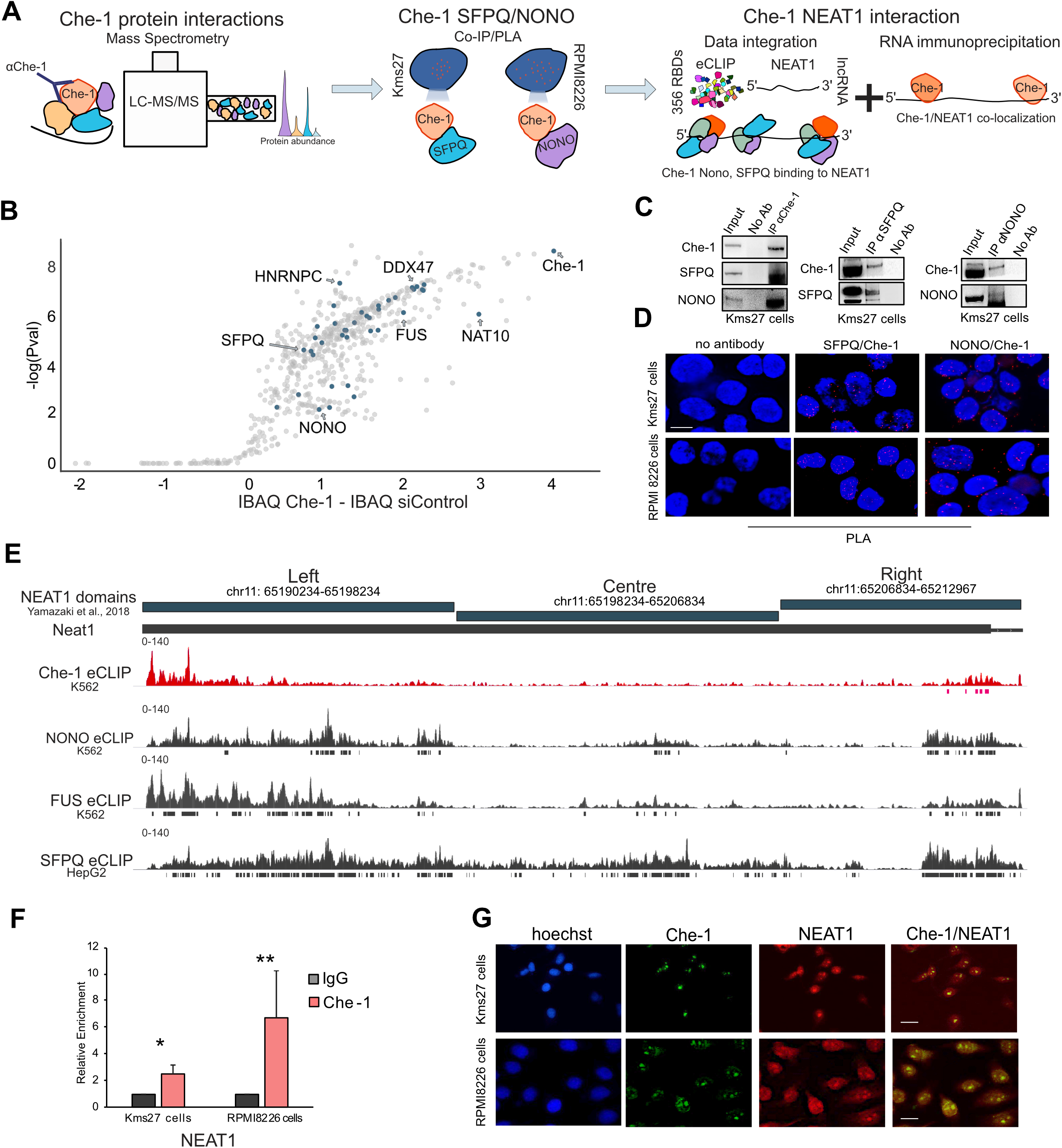
Che-1 interacts with paraspeckle components. **A:** Experimental and computational workflow followed to test the relationship between Che-1 and paraspeckle components. *(Left)* Mass spectrometry analysis was performed to identify Che-1 interactome followed by: *(Middle)* Che-1 SFPQ/NONO co-immunoprecipitation (Co-IP) and Proximity ligation assay (PLA) to evaluate physical interaction. (*Right)* Then eCLIP data of 356 RNA Binding Protein in K562 cell line was evaluated at NEAT1 locus followed by Che-1 RNA immunoprecipitation (RIP) and co-localization in KMS27 and RPMI8226 Multiple Myeloma (MM) cell lines. **B:** Scatter Plot of mass spectrometry assay. Dots depict protein intensity (IBAQ) and relative significance obtained from mass spectrometry assay. X-axis: Difference of Che-1 intensity versus IP control; on the y-axis are the corresponding −log10 P-values. Significant Che-1 interacting proteins relevant in this study are represented in blue. For a complete list of all significant Che-1 interactors, please refer to Supplementary Table S1. **C:** Total extract from Kms27 MM cells was subjected to immunoprecipitation with the indicated antibodies. Negative control was performed with IgG. Input corresponds to 5% of the total extract used for immunoprecipitation. One out of two experiments is shown. **D:** Representative images of PLA showing the interaction between Che-1 and paraspeckles protein SFPQ or NONO in Kms27 and RPMI8226 MM cells. Negative controls were performed by omitting Che-1 primary antibody whilst nuclei were stained with Hoechst dye. One out of two experiments for each line is shown. Scale bar 10μm. **E**: eCLIP binding enrichment of Che-1, NONO, FUS, SFPQ at NEAT1 domains. NEAT1 domains were defined in Yamazaki et al.(46), and are represented as blue domains (Top). Genomic coordinates represent the position of each NEAT1 domain in the hg19 reference genome. eCLIP profiles were obtained from Van Nostrand, et al.(45), and plotted at a fixed scale (0–140) using IGV. Replicates of each experiment were merged in a unique overlayed profile for each experiment. Che-1 (Red), NONO (Grey), FUS (Grey) eCLIP profiles were obtained from K562 cells while SFPQ eCLIP (Bottom) from HepG2 cells due to the relative K562 cells experiment unavailability. **F:** RIP assay performed in Kms27 and RPMI8226 MM cells using an anti-Che-1 specific antibody. IgG were used as negative control. NEAT1 RNA abundance was evaluated by RT–qPCR. Results from at least three biological replicates for each line are shown. Statistical significance is indicated by asterisks as follows: \**P*< 0.018. Statistical analysis was performed using two-sided t-tests. **G:** Colocalization assays were performed in Kms27 and RPMI8226 MM cell lines by combining FISH staining of NEAT1 (Stellaris) with immunofluorescence of Che-1. Nuclei were stained with Hoechst dye. The experiment was performed two times, obtaining similar results. Representative images for each cell line are shown. Scale bar 10μm.

Overall, our data support the hypothesis that Che-1 is a component of paraspeckles by interacting directly with NEAT1.

### Che-1 and NEAT1 colocalize on the DNA

Previous studies identified the lncRNA NEAT1 as an essential structural determinant of paraspeckles. It has been demonstrated that these structures regulate gene transcription in response to different types of stress by sequestrating specific RNAs and proteins within them (15), (47). Moreover, NEAT1 is detectable bound at the DNA of nucleosome depleted regions associated to active transcription sites (18). To investigate the potential co-binding affinity of NEAT1 and Che-1 at DNA we firstly identified NEAT1 binding site with chromatin isolation by RNA purification (ChIRP-seq) and Che-1 binding with chromatin immunoprecipitation (ChIP-seq) experiments in Kms27 MM cells and developed computational signal dissection approach to define evidence of co-binding at same loci. Our approach relied on the identification at high resolution of NEAT1 and Che- 1 binding sites followed by a quantification of both signals at potential ∼92k Transcription Starting Sites (TSS). ChIRP- seq and ChIP-seq signals were independently normalized, ranked for Jaccard similarity index. Finally, signals we assigned to 3 groups accordingly to mathematical evidence: NEAT1/Che-1 co-binding, only Che-1 and only NEAT1 (Figure 2A).

**Figure 2:**
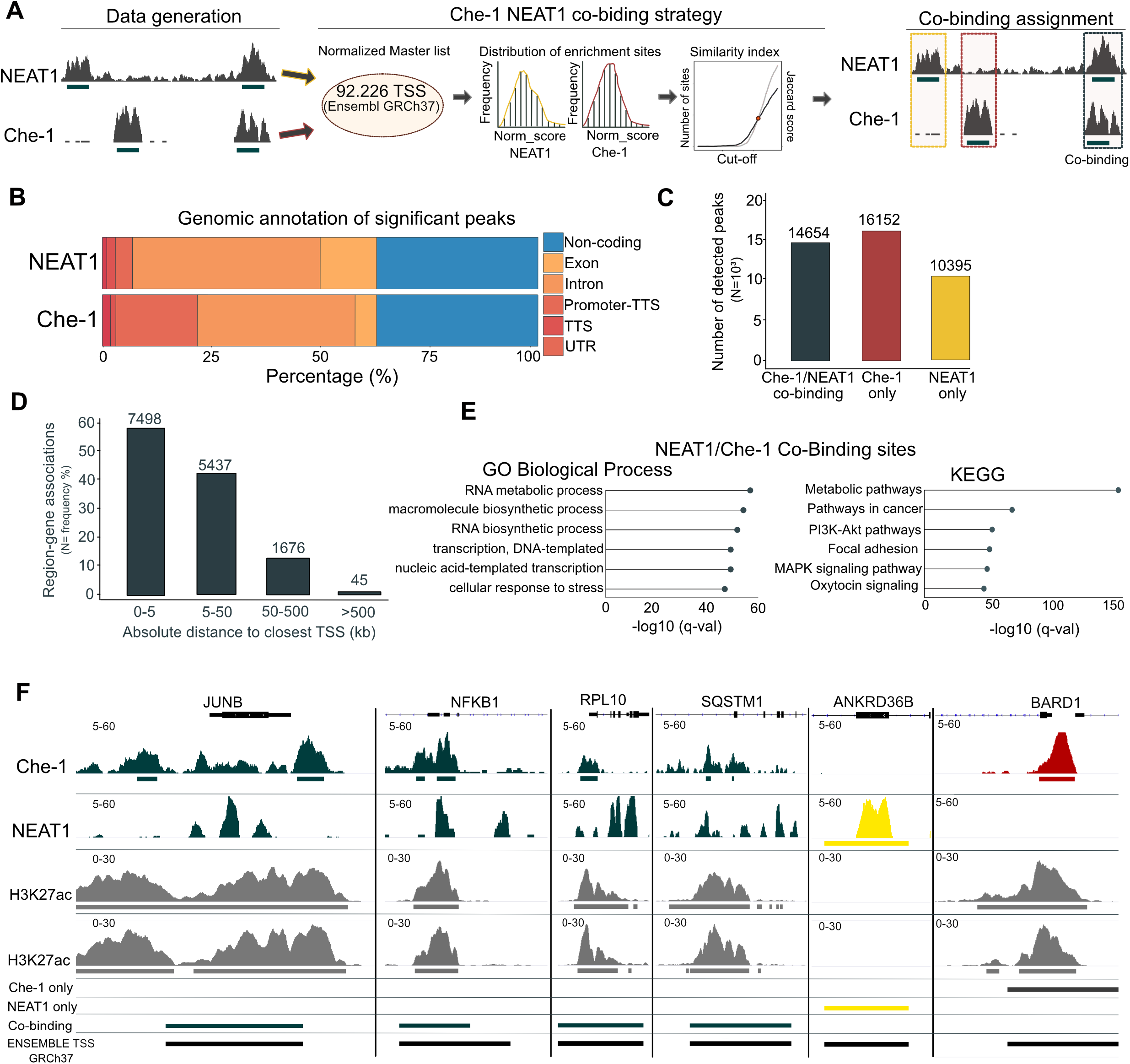
Che-1 and NEAT1 colocalize on the DNA. **A:** Experimental and computational workflows developed to identify NEAT1/Che-1 co-binding sites in MM Kms27 cells. ChIRP-seq and ChIP-seq were used to identify NEAT1 and Che-1 biding sites, respectively. Then, normalized read enrichment of both experiments was assessed at 92,226 transcription starting sites (coordinates from Ensembl GRCh37). Peaks were classified using an iterative approach of selection and assignment based on Jaccard scoring (Middle). Finally, peaks were classified into only NEAT1, only Che, Che-1/NEAT1 co-binding. **B:** Genomic annotation of significant peaks from the NEAT1 ChIRP-seq (top bar) and Che-1 ChIP-seq (bottom bar) experiments. Peaks were classified among six categories: Non-coding, Exon, Intron, Promoter-TTS, TTS, UTR according to Homer annotation tool. Each category is shown with a different color. The percentage of classified peaks for each category is shown as experiment-specific percentage (x-axis). **C:** Boxplot showing the number of peaks shared between NEAT1/Che-1 (dark green), Che-1 only, (red), NEAT1 only (yellow). The peak number is shown at the top of each bar. **D:** Boxplot showing the relative distance of each Che-1/NEAT1 co-binding peak to the closest gene. Distance is calculated as absolute value and is expressed in kilobase. The number of peaks per category at the top of each bar. Bar height represents the proportion of peaks included in each category to the total (N=14654). **E:** Gene Ontology (left) and KEGG enrichment pathways analysis (right) of Che-1/NEAT1 co- binding sites. The top 6 enriched terms are shown. Only ontology terms significant by both the binomial and hypergeometric tests using the multiple hypothesis correction false discovery rate are considered in this analysis. The represented q-value (x-axis) corresponds to the hypergeometric test.

ChIRP-seq enriched at 19,1369 regions while Che-1 ChIP-seq at 41,061 binding sites (Supplementary Figure S2a). Enriched regions were classified according to their genomic position. Che-1 was more prone to bind downstream the promoter (TTS, -100 bp to +1kb genomic window) while NEAT1 at gene bodies (Figure 2B). Interestingly, ∼30% of Che-1 and NEAT1 sites bound at non-coding regions thus suggesting a functional role for those sites which could be further investigated. These data agree with NEAT1 and Che-1 being key determinants of gene transcription regulation. To gain insights into the binding at the TSSs we leveraged the genomic annotation obtained from Ensembl GRCh37 by surveying the signal at 92,226 sites. We identified those sites showing Che-1/NEAT1 co-binding (N=14,654), Che-1 only sites (N=16,152) and NEAT1 only sites (N=10,395) (Figure 2C). Most co-binding sites (87% of the totality) were occurring at promoter proximity (Distance to promoter TSS between 0-50 Kb) (Figure 2D). Ontologies and pathways analysis for the genes associated to those sites showed highly significant enrichments for active transcription and metabolic process (Figure 2E). These data prompted us to investigate whether Che- 1/NEAT1 co-binding was occurred at open chromatin sites. Accordingly, we profiled H3K27ac ChIP-seq data of Kms27 cell lines and integrated the signal with NEAT1 and Che-1 sites (Figure 2F). Strikingly 47% of co-binding sites was overlapping with H3K27ac marked regions while Che-1 only and NEAT1 only sites were occurring at active sites respectively at 36% and 11%, respectively (Supplementary Figure S2B). Taken together, these data strongly support a potential regulatory role for Che-1 in occurrence with NEAT1 at active genes.

### NEAT1 is required for recruiting Che-1 onto the DNA

Next, we investigated whether the Che-1/NEAT1 interaction could affect the localization of Che-1 in the nucleus. As shown in Figure 3A, NEAT1 depletion by specific Antisense LNA GapmeR in Kms27 MM cells produced a dispersion of Che-1, which assumed a perinuclear localization. Similar results were obtained in another MM cell line RPMI8226 (Supplementary Figure S3A). To confirm these results, we probed the presence of Che-1 in the various cellular components. As shown in Figure 3B, the downregulation of NEAT1 produced an accumulation of Che-1 in the nucleoplasm, together with a concomitant decrease in the histone H3 acetylated and total chromatin fraction. Consistent with these findings, Che-1 ChIP-seq analysis revealed a marked global decrease of Che-1 occupancy at the Che-1/NEAT1 colocalization sites in NEAT1 depleted cells (Figure 3C). Remarkably, NEAT1 depletion in MM cells recapitulated the effects of Che-1 downregulation (7), with a reduction in cell proliferation and global RNA synthesis and a significant downregulation of several genes involved in MM pathogenesis, such as *IRF4* or *MMSET* (*7*) (Supplementary Figures S3B, S3C and S3D) . To further characterize the Che-1/NEAT1 interaction, we evaluated the effects of Che-1 downregulation on the presence of NEAT1 onto the DNA. Fluorescence In Situ Hybridization (FISH) analysis performed in Kms27 cells showed that Che-1 depletion does not affect NEAT1 localization (Figure 3D). Moreover, ChIRP-seq analysis performed in Che-1 depleted Kms27 cells showed that only a marginal fraction of NEAT1 peaks were affected by Che-1 depletion (Figures 3E and 3F). Indeed, approximately 90% of Che-1 and NEAT1 colocalizing sites, maintained similar NEAT1 enrichment in response to Che-1 downregulation. Taken together, our results show that NEAT1 is required for recruiting Che-1 onto the DNA, whereas NEAT1 binding on DNA is not dependent to Che-1 occurrence.

**Figure 3:**
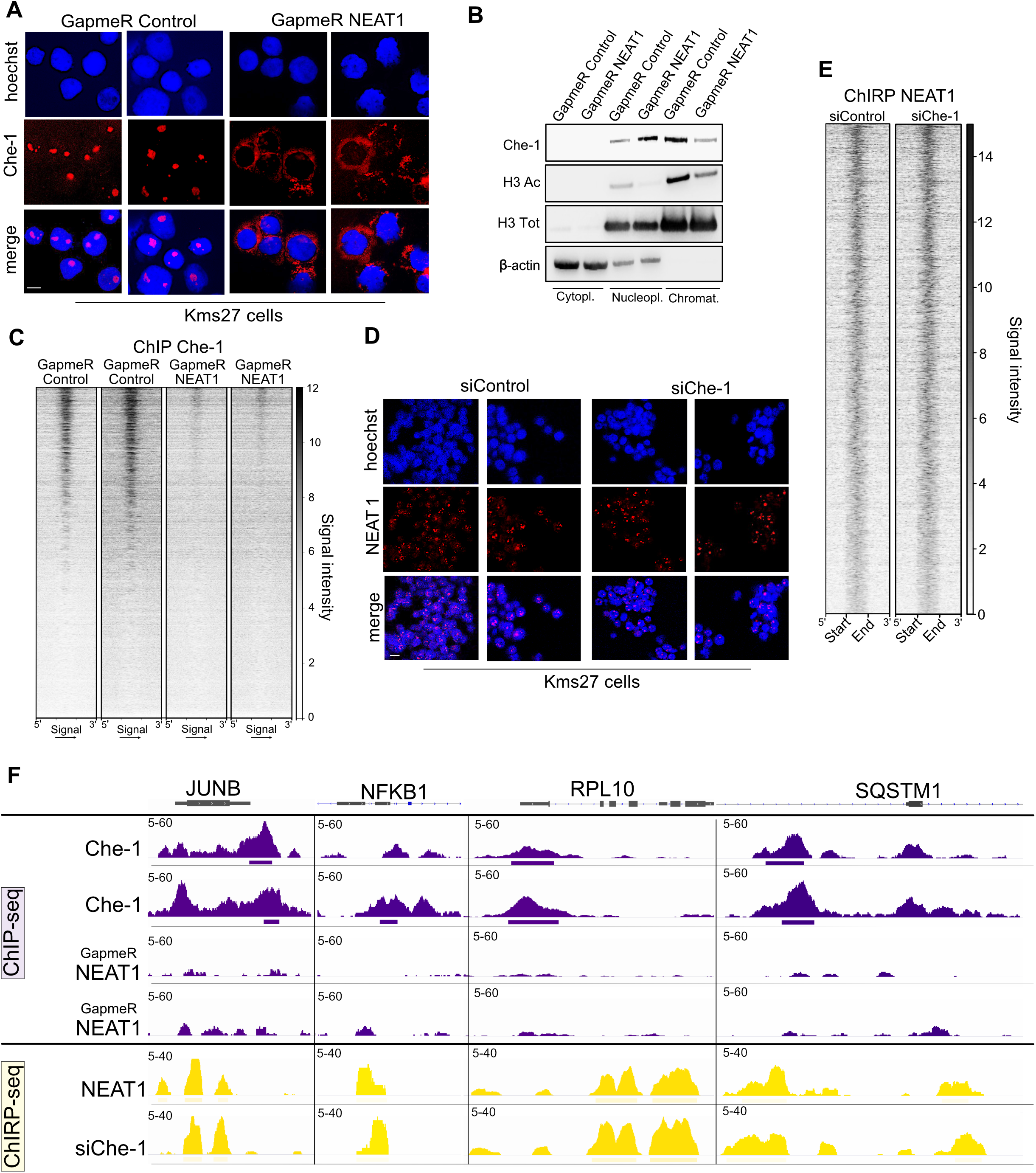
NEAT1 is required for recruiting Che-1 onto the DNA. **A:** Representative immunofluorescence images of Kms27 MM cells fixed and immunostained with anti-Che-1 antibody after transfection with NEAT1 or control Antisense LNA GapmeR oligonucleotides. One of two experiments is shown. Scale bar 10 μm. **B:** Isolated cell compartments from Kms27 MM cells transfected with GapmeR Control or GapmeR NEAT1 oligonucleotides, were subjected to western blot analysis with the indicated antibodies. Data shown represent two independent experiments. **C:** Che-1 ChIP-seq signal intensity of GapmeR Control (2 replicates on left) and GapmeR NEAT1 (2 replicates on right) at co-localizing Che-1/NEAT1 sites (N=14654 sites). Signal intensity from 1 (weak) to 12 (strongest). **D:** Fluorescence In Situ Hybridization (FISH) images of Kms27 MM cells depleted or not for Che-1 expression labeled with Hoechst (cell nuclei) and Quasar 670 (NEAT1 RNA). Scale bar 10 μm. **E:** NEAT1 ChIRP-seq signal intensity of Kms27 MM cells transfected with siControl or siChe-1 at co-localizing Che-1/NEAT1 sites (N=14654 sites). Signal intensity from 1 (weak) to 14 (strongest). **F:** Che-1 ChIP-seq signal (top, in violet) in GapmeR Control (top, 2 replicates) and GapmeR NEAT1 (bottom, 2 replicates) and NEAT1 ChIRP-seq (yellow) of siControl control (top) and siChe-1 (bottom) at JUNB, NFKB1, RPL10, SQSTM1 loci. Signal enrichment scale on the y-axis of each site.

### Che-1 and NEAT1 suppress RNA:DNA hybrids accumulation

While these analyses supported a strong interplay between NEAT1 and Che-1, the functional significance of this interaction was unknown. Recently, a mass spectrometry study identified the R-loops interactome in human cells, dynamic structures which contribute to gene regulation in health and disease (48). Interestingly, RNA:DNA hybrids and Che-1 interactomes shared a large percentage of proteins suggesting that Che-1 and paraspeckles could be present on the RNA:DNA hybrids in MM cells (Figure 4A). To this aim, co-immunoprecipitation experiments were performed with Kms27 and RPMI8226 cell lysates by using the S9.6 antibody, which recognizes RNA:DNA hybrids (49). These experiments showed that Che-1, NONO and SFPQ bind RNA:DNA hybrids, and that these interactions are specific, since they disappeared after RNase H treatment (Figure 4B). Consistent with these results, PLA experiments revealed the presence of Che-1 on RNA:DNA hybrids (Figure 4C). To assess whether NEAT1 was also present on RNA:DNA hybrids, we performed RNA immunoprecipitation (RIP) experiments using the S9.6 antibody. As shown in Supplementary Figure S4A, NEAT1 was found on these hybrids in both Kms27 and RPMI8226 MM cells. Notably, the presence of Che-1, SFPQ and NONO on the RNA:DNA hybrids appeared to be mediated by NEAT1, since the depletion of this lncRNA strongly reduced the interaction between these three proteins and RNA:DNA hybrids (Figure 4D). To investigate the role of Che-1 in R-loops metabolism, we depleted Che-1 expression in Kms27 cells by specific siRNA oligonucleotides. As shown in Figures 4E and 4F, Che-1 down-regulation produced a strong accumulation of R-loops in these cells which disappeared after treatment by RNase H. Of note, NEAT1 depletion also induced an accumulation of R-loops in Kms27 cells (Supplementary Figure S4B). These findings were confirmed by performing specific DRIP-qPCR of genes with a high frequency of R-loop s(50). Indeed, Che-1 depletion induced a strong increase of RNA:DNA hybrids on these regions, (Supplementary Figure S4C). Finally, RNA:DNA immunoprecipitation and high-throughput sequencing (DRIP-seq) demonstrated that Che-1 depletion causes an accumulation of cellular R-loops on the Che-1/NEAT1 co-binding sites (Figures 4G and 4H). Taken together, these results demonstrate that Che-1 and NEAT1 cooperates in preventing R-loops accumulation.

**Figure 4:**
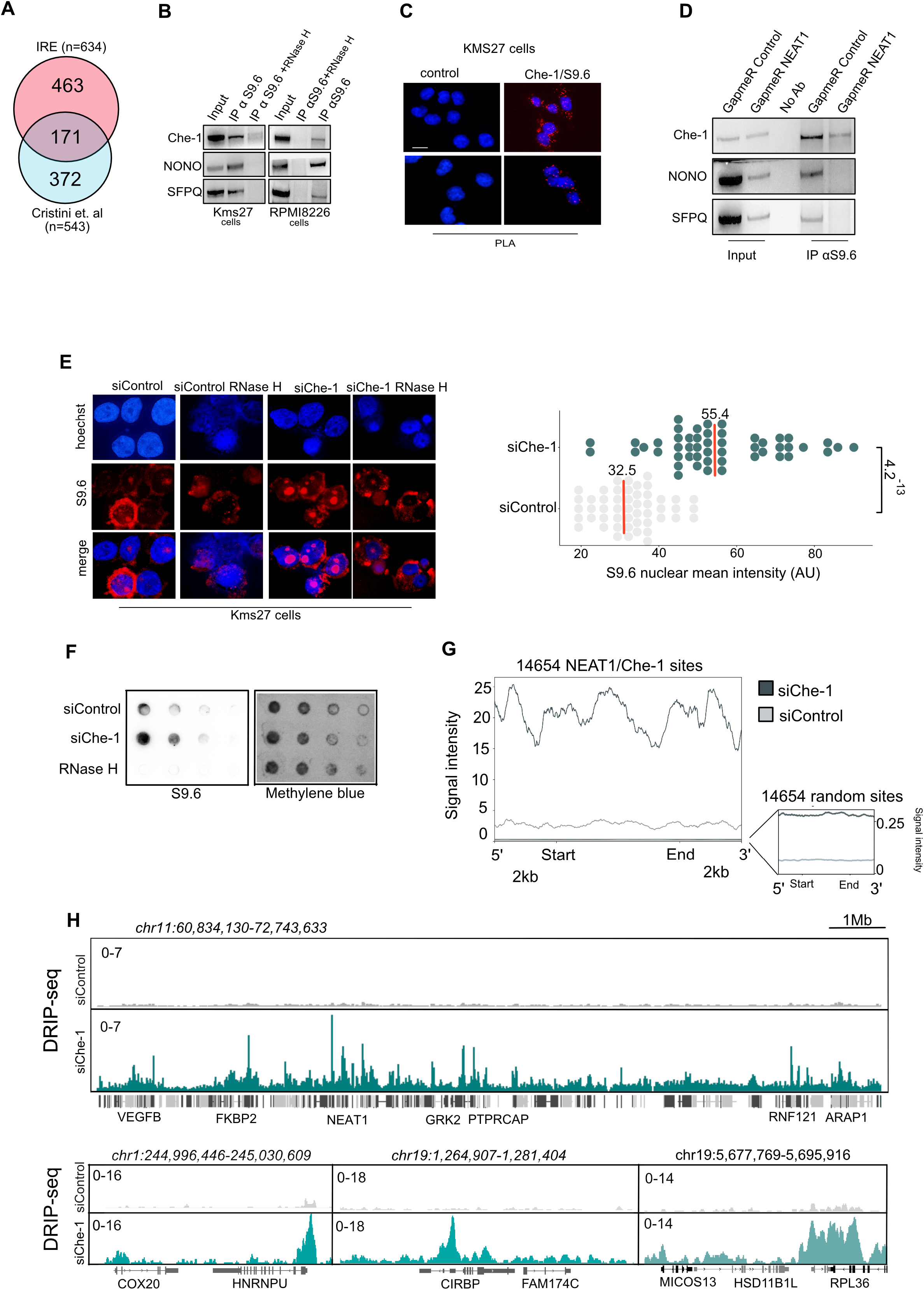
Che-1 and NEAT1 suppress RNA: DNA hybrids accumulation. **A:** Venn diagram showing the relationship between enriched protein in Che-1 and RNA:DNA hybrids interactome respectively obtained from this study and Cristini et al.(48) study. **B**: Total extracts from Kms27 and RPMI8226 MM cells treated or not with RNase H, were subjected to immunoprecipitation with S9.6 antibody. Immuno-precipitated complexes were then analyzed by western blot with the indicated antibodies. Input corresponds to 5% of the total extract used for immunoprecipitation. One out of two experiments is shown. **C:** Representative images of Kms27 MM cells showing the interaction between Che-1 and S9.6 revealed by PLA. Negative controls were performed by omitting Che-1 primary antibody, whilst nuclei were stained with Hoechst dye. Scale bar 10 μM. **D:** Total extracts from Kms27 MM cells transfected with LNA GapmeR Control or GapmeR NEAT1 oligonucleotides were immunoprecipitated with S9.6 antibody. Immuno- precipitated complexes were then analyzed by western blot with the indicated antibodies. Input corresponds to 5% of the total extract used for immunoprecipitation. One out of two experiments is shown. **E:** (*left*) Immunostaining of Kms27 MM cells with S9.6 antibody (red) and Hoechst (blue), transfected with siControl or siChe-1 and treated or not with RNase H. Scale bar 10 μM. (*right*) Dot plot showing S9.6 signal nuclear intensity per cell (Arbitrary Unit-AU). siChe-1 cells are described in green color, while siControl in grey color. 100 cells were counted in each replicate. Each dot represents a cell. Median scores per sample are written at the top of the median red line. *P* value: 4.2^-13^. T-test performed to evaluate statistical significance. **F**: Dot blot analysis of RNA: DNA hybrids formation by serial dilutions of genomic DNA starting at 1.5 micrograms were spotted on a membrane and probed using the S9.6 antibody. Methylene blue normalization of S9.6 signal image was acquired by using Alliance Mini HD6 system by UVITEC Ltd,. One out of two experiments are shown. **G:** Enrichment density of DRIP-seq signals at NEAT1/Che1 co-localizing sites in Kms27 MM cells identified in siRNA control (siControl) (grey) and siRNA Che-1 (siChe-1) (dark green) samples. Enrichment density was calculated on the relative size of each peak (start to end of each fragment) extended 2000 bp upstream and downstream. Right frame represents the signal obtained from 14654 randomly selected sites from the same experiments. **H:** DRIP-seq signal in siControl (grey) and siChe-1 (green) samples at NEAT1 genomic window (top) and 3 genomic windows (bottom). Genomic coordinates described above the signals. Signal enrichment scale on the y-axis at each site.

### Che-1 controls IFN activation in MM cells

The results shown above demonstrate how Che-1 interacts with NEAT1 on DNA and contributes to the control of RNA:DNA hybrids formation. Che-1 depletion is indeed sufficient to induce increase of RNA:DNA hybrids. Consistent with these findings, RNA sequencing experiments (RNA-seq) in Kms27 cells revealed a strong upregulation of many genes involved in NF-kB and both interferon (IFN) type I (IFN alpha and beta) and type II (IFN gamma) signaling in Che-1 depleted cells (Figures 5A, 5B and 5C), in line with what observed in numerous cell models where antiviral response is a fundamental mechanism in response to DNA damage (51, 52). Indeed, Che-1 depleted cells exhibited a strong activation of the RNA:DNA sensor pathway cGAS-Sting (53, 54) and the RNA helicase RIG-1, involved in double-stranded RNA-induced innate antiviral response (55) (Figure 5D). Notably, similar results were obtained from an RNA-seq analysis of Kms27 cells with or without NEAT1 depletion (Supplementary Figures S5A, S5B and S5C), underscoring a common functional role played by Che-1 and NEAT1. We validated these results by taking advantage of a NF- kB-luciferase reporter assay. As shown in Figure 5E, Che-1 depletion was able to activate luciferase activity at the same levels observed by the overexpression of RelA protein, the principal member of the NF-kB complex (56). Of note, both IFNB1(Interferon beta1) and IFNG (Interferon gamma) were upregulated in Che-1 depleted cells (Figure 5F), and the increase of transcript abundance of NF-kB and IFN target genes confirmed the activation of these pathways (Figure 5G). In agreement, we found similar results in NEAT1 depleted Kms27 cells (Supplementary Figures S5D and S5E). Strikingly, IFN activation induced by Che-1 depletion was completely reversed by overexpressing GFP-RNase H (Figures 5H and 5I). Overall, these results demonstrate that Che-1 and NEAT1 prevent IFN response in MM cells by facilitating RNA:DNA hybrids resolution.

**Figure 5:**
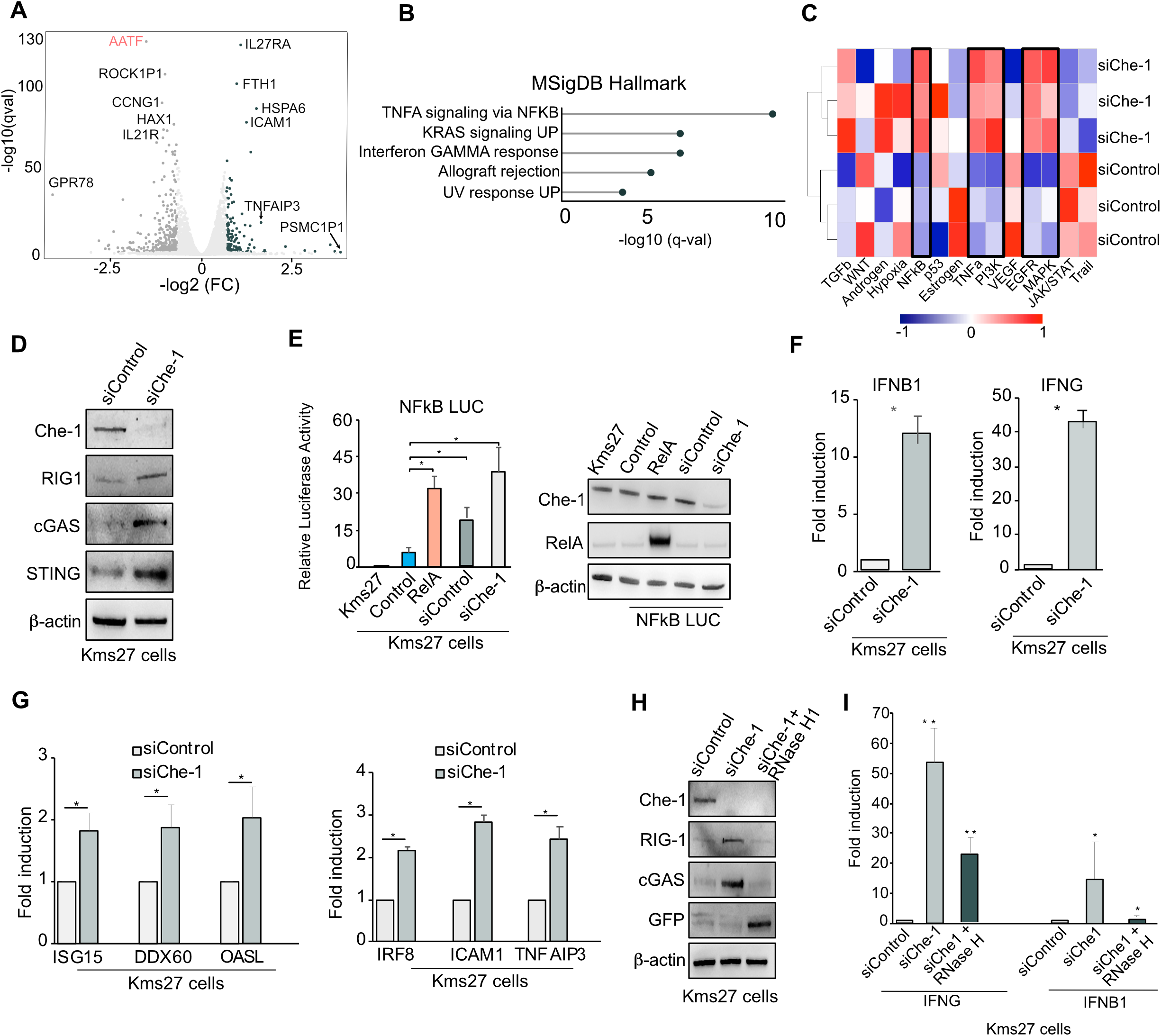
Che-1 controls Interferon activation in MM cells. **A:** Differential analysis of siControl vs. siChe1 transcriptome in Kms27 MM cells. Volcano plot shows 135 significantly upregulated (dark green) and 392 downregulated genes (dark grey). x-axis reports the base-2 logarithm of fold change which is approximated to the B value coming from the statistical Wald test. y-axis reports the base-10 logarithm of Q value of significant genes. **B:** Gene Set Enrichment Analysis (GSEA) of upregulated significant differential genes with their relative base-10 logarithm of Q value. The analysis was based on hallmark gene sets collection. **C:** Heatmap depicting pathway activity scores for each transcriptome sample according to PROGENy. The activity score is shown as shade ranging from blue to red which indicates the level of pathway activation. Blu= non-active; Red=active. **D:** Western blot analysis of total extracts of Kms27 MM cells depleted or not for Che-1 and probed for the indicated antibodies. Data shown represent two independent experiments. **E:** *(left)* Luciferase activity of human NF-kB promoter was measured in Kms27 MM cells transfected with siChe-1 or siControl oligonucleotides or with empty vector or RelA protein expressing vector. All data are expressed as a percentage of control value and presented as mean ± SD of three independent experiments. Statistical significance is indicated by asterisks as follows: \**P*< 0.004. Statistical analysis was performed using two-sided t-tests. *(right)*Western blot analysis of total extracts of Kms27 MM cells transfected as described above and incubated with indicated antibodies. **F and G:** RT-qPCR analysis of the levels of the indicated genes in Kms27 MM cells depleted or not for Che-1. Data are presented as mean ± SD of three independent experiments. Statistical significance is indicated by asterisks as follows: \**P*< 0.0075. Statistical analysis was performed using two-sided t-tests. **H:** Western blot analysis of total extracts of Kms27 MM cells transfected with siChe-1 or siControl and with GFP-tagged RNaseH and incubated with indicated antibodies. Data shown represent two independent experiments. **I:** RT–qPCR analysis of IFNB1 and IFNG levels in Kms27 MM cells depleted or not for Che-1 and treated or not where indicated with RNase H. Statistical significance is indicated by asterisks as follows: \**P*< 0.04, \*\**P*< 0.004. Statistical analysis was performed using two-sided t-tests.

### MM patients exhibit high levels of RNA:DNA hybrids

These analyses support a pivotal role of the Che-1/NEAT1 interaction in conferring a survival advantage to cells during stress. We first translated these findings by leverage ∼1100 MM patient transcriptomes from the CoMMpass data set and Oncomine database (57), then we evaluated RNA:DNA hybrids levels in a MM patient’s cohort. Previous studies have demonstrated a consistent association between the expression of Che-1 and NEAT1 and the progression of MM (6,7,58,59). The analysis of Oncomine database (57) confirmed these observations (Supplementary Figure S6A). This prompted us to investigate whether same results would have been obtained from querying the International Staging System (ISS) available in the CoMMpass dataset. Notably, this analysis showed that not only the expression of Che-1, but also of NEAT1 and numerous components of the paraspeckle, were significantly higher in ISS stage 3 compared to ISS 1 and ISS 2 (Supplementary Figure S6B). In addition, survival analysis of 542 MM (60) showed that high expression of Che-1 and NEAT1 are strongly associated with poorer prognosis in MM (Supplementary Figure S6C). Then we assessed RNA:DNA hybrids levels in MM patients. Remarkably, immunofluorescence analysis of 15 samples of CD138^+^ MM cells and 5 samples from healthy controls showed a significant higher level of hybrids in most MM patients (Figures 6A, 6B and 6C), and both cGAS and RIG-1 levels strongly increased in MM cells compared with normal plasma cells (Figure 6D). Accordingly, transcriptomic analysis if CoMMpass dataset revealed a close association between ISS grade and activation of IFN pathway (Figures 6E and 6F). Moreover, a high expression of this pathway was associated with poorer prognosis in the patient cohort selected by IFN gene signature expression (Figure 6G, Table 1 and Supplemental Table S4) and in the selection of patients exhibiting the highest expression of the signature (Figure 6H). Taken together, these results indicate a strong association between elevated levels of RNA:DNA hybrids, the expression of paraspeckle genes, and the activation of the IFN response.

**Figure 6:**
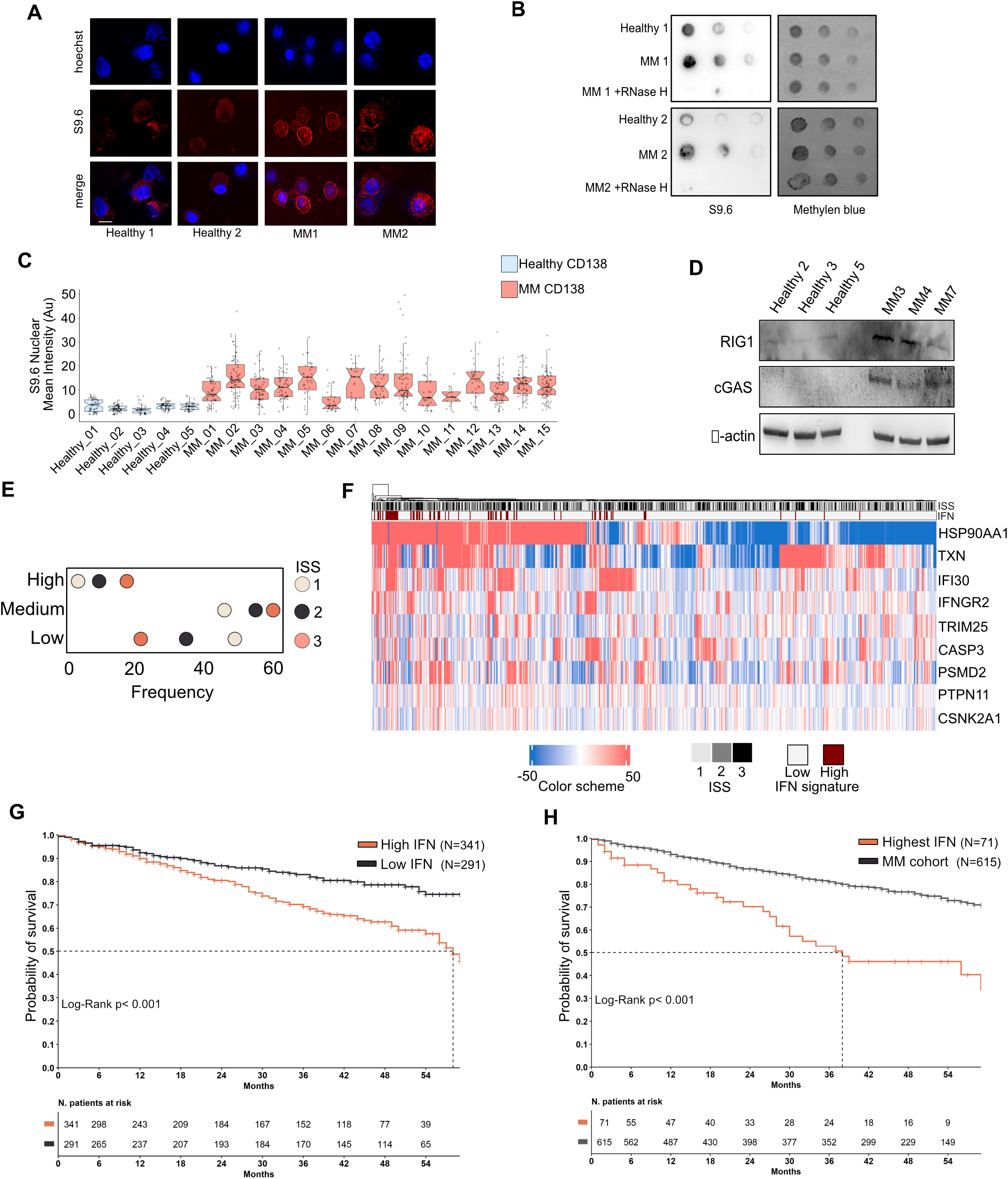
MM patients exhibit high levels of RNA: DNA hybrids. **A:** Representative S9.6 immunofluorescence levels in healthy and tumor CD138^+^ plasma cells purified from patient bone marrow. Nuclei were visualized by staining with Hoechst dye. Scale bar 10μm. **B:** Dot blot performed to evaluate the enrichment of RNA:DNA hybrids formation using the S9.6 antibody by serial dilutions of genomic DNA starting at 1.0 micrograms. Methylene blue normalization of S9.6 signal image acquired by using Alliance Mini HD6 system by UVITEC Ltd,. One out of two experiments are shown. **C:** Boxplot showing relative S9.6 nuclear mean intensity in CD138^+^ plasma cells from 15 MM patients (red) and 5 healthy (light blue). Dot plot representing each cell considered overlay the boxplot. Y-axis: S9.6 Nuclear Mean Intensity (AU). **D:** Representative WB analysis with the indicated antibodies of healthy and neoplastic CD138^+^ purified plasma cells from patients. **E:** Proportion of CoMMpass patients (N=687) assigned to the IFN signature category in relationship with the relative ISS stage. Each patient was assigned to the IFN signature category: High (top), Medium (middle) Low (bottom). Dot colors correspond to the ISS stage of samples for each IFN signature category. **F:** Unsupervised clustering heatmap showing enrichment profiles of 9 IFN signature genes. From the top: ISS stages 1,2,3 of each patient (N=687). IFN signature High (red, N=71) and Low (white, N=614) assigned to each patient. Color scheme representing up- (red) and down- (blue) regulation of the selected genes across the patient dataset. Colors are proportional to normalized (cpm) read enrichment – the relative read enrichment median of each gene within the full patient cohort. **G:** Kaplan–Meier survival curves of overall survival (OS) in the CoMMpass cohort (N = 687) with IFN signature selection expression on a continuous scale. The patient cohort was divided in IFN high (N=341) (red) and IFN low (N=291) (black) by splitting the population at the IFN signature median threshold. The patients above the IFN median were assigned to IFN High, while patients below the median were assigned to IFN low group. **H:** Kaplan–Meier survival curves of overall survival (OS) in theCoMMpass cohort (N = 687) with IFN signature selection on a discrete scale. The patient cohort was sp lit in IFN highest (N=71) (red) and the remaining MM cohort (N=615) (black) as described in the methods section.

### UPR induced R-loops in MM cells

We then sought to identify the mechanism underlying the elevated levels of RNA:DNA hybrids that we found in MM. Recently, it has been demonstrated that the Integrated Stress Response (ISR) induces R-loops in U2OS cells (61). Since a peculiar characteristic of MM cells is a high Unfolded Protein Response (UPR) due to their abnormal production of immunoglobulins (62), it is conceivable that this type of stress may contribute to the generation of R-loops. To test this hypothesis, we took advantage of two different MM cell lines endowed with different levels of UPR (Kms27 high UPR, U266 low UPR). As shown in Figures 7A, 7B, 7C and Supplementary Figure S7B, elevated UPR activation in Kms27 cells was found strongly associated with higher levels of R-loops and IFN expression when compared to U266 cells. In agreement, both Che-1 and NEAT1 also resulted more expressed in Kms27 than U266 cells (Figure 7A and Supplementary Figure S7A). To further investigate this association, we induced UPR in U266 cells using thapsigargin (THAP). Increased expression of ATF4, PERK and IRE1a confirmed UPR activation, together with an augment of Che-1 and NEAT1 levels (Figure 7D and Supplementary Figure S7C). Strikingly, treatment with THAP produced a dramatic increase of both RNA:DNA hybrids and IFN expression (Figures 7E, 7F and Supplementary Figure S7D). To confirm these findings, we treated Kms27 cells with ISRIB, a selective inhibitor of Integrated Stress Response (ISR) that impairs adaptation to ER stress (63–65). As shown in Figure 7G, Kms27 cells treated with this drug exhibited a strong reduction of UPR activation, and both Che-1 and NEAT1 expression, associated with decreased levels of R-loops and IFN activity (Figures 7G, 7H, 7I, Supplementary Figures S7E and S7F). Overall, these results demonstrate that UPR induces R-loops and inflammatory signaling in MM cells.

**Figure 7:**
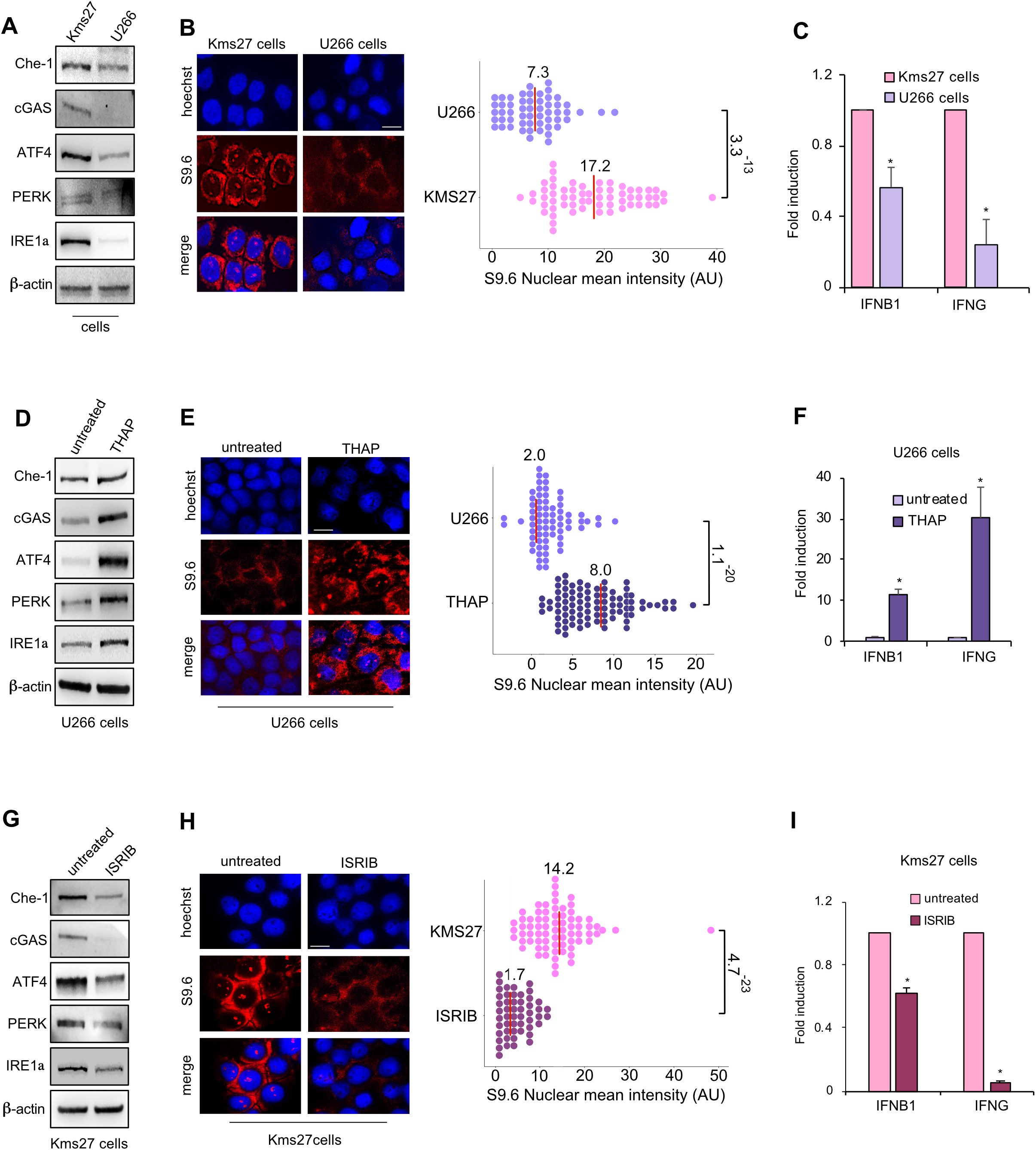
UPR induced R-loops in MM cells. **A:** Western blot analysis of total extracts from Kms27 and U266 human MM cells with the indicated antibodies. **B:** (*left*) Kms27 and U266 MM cells were subjected to immunostaining with S9.6 antibody (red) and Hoechst (blue). One out of two experiments is shown. (*right*) Box plot showing relative S9.6 nuclear mean intensity cell in the nucleus. 100 cells were counted in each replicate. Median scores per sample are written at the top of the median red line. *P* value: 3.3^-13^. T-test performed to evaluate statistical significance. **C:** Different expression of IFNG and IFNB1 in Kms27 and U266 MM cells was assessed by RT-qPCR. Results from three biological replicates are shown. Statistical significance is indicated by asterisks as follows: \**P*< 0.002. Statistical analysis was performed using two-sided t-tests. **D:** Representative WB analysis with the indicated antibodies of total extracts from U266 MM cells treated or not with Thapsigargin (THAP) 0.5 μg/ml for 16 hours. **E:** (*left*) Immunostaining with S9.6 antibody in U266 MM cells treated or not with THAP as in **D.** One out of two experiments is shown. (*right*) Box plot showing relative S9.6 nuclear mean intensity per cell. 100 cells were counted in each replicate. Median scores per sample are written at the top of the median red line. *P* value: 1.1^-20^. T-test performed to evaluate statistical significance. **F:** RT-qPCR analysis of IFNG and IFNB1 in U266 MM cells treated or not with THAP. Results from three biological replicates are shown. Statistical significance is indicated by asterisks as follows: \**P*< 0.0025. Statistical analysis was performed using two-sided t-tests. **G:** Immunoblot with indicated antibodies of total extracts from Kms27 MM cells treated or not with ISRIB 1μM for 6 hours. **H:**(*left*) Immunostaining with S9.6 antibody of Kms27 MM cells treated or not with ISRIB as in **G.** One out of two experiments are shown. (*right*) Box plot showing relative S9.6 nuclear mean intensity per cell. 100 cells were counted in each replicate Median scores per sample are written at the top of the median red line. *P* value: 4.7^-23^. T-test performed to evaluate statistical significance. **I:** RT-qPCR analysis of IFNG and IFNB1 levels in Kms27 MM cells treated as in **G**. Results from three biological replicates are shown. Statistical significance is indicated by asterisks as follows: \**P*< 3.27^-05^. Statistical analysis was performed using two-sided t-tests. Data shown in **A**, **D** and **F** were performed two times with similar results obtained.

## Discussion

In this study we demonstrate that Che-1 is a component of paraspeckles. Che-1 physically interacts with the NEAT1 lncRNA in MM cells and provide evidence that these two molecules widely colocalize on DNA sites. We show that Che-1 and NEAT1 bind R-loops and are required for their resolution, preventing their accumulation and inflammatory response activation. We also show that high levels of RNA:DNA hybrids are present in MM patients and that activation of IFN response correlates with the stage of the disease and is a negative prognostic factor. Finally, we provide evidence that unfolded protein response (UPR) in MM cells promotes the formation of R-loops and the consequent activation of IFN response.

Paraspeckles are subnuclear bodies of which the NEAT1 lncRNA is the main structural component, necessary for their formation and maintenance (14). Although numerous studies over the last few years have outlined a role of these structures in RNA and protein retention, NEAT1 has also been found to localize to transcriptional start sites of numerous genes, hypothesizing its active role in the regulation of gene transcription (18). Our results confirm the presence of NEAT1 on DNA in over 25,000 sites (Figure 2C), with an accumulation on the coding regions, reinforcing the finding that NEAT1 plays an important role in transcription. Moreover, our results demonstrate that NEAT1 and Che-1 colocalize onto the DNA, and that NEAT1 is required for recruiting Che-1 on these sites. Therefore, it is conceivable a scenario in which NEAT1 can act as a hub to concentrate paraspeckles components involved in the transcription or maturation of RNA on specific DNA sites. Our results also demonstrated that Che-1 and NEAT1 interact with the RNA:DNA hybrids present on DNA, and that the presence of NEAT1 is required to recruit not only Che-1, but also other paraspeckle components such as NONO or SFPQ onto these structures, reinforcing the hypothesis that NEAT1 performs an important hub function. Of note, NEAT1, and especially Che-1, appear necessary for the correct resolution of R-loops, given the massive accumulation of these structures on the binding sites of Che-1, when the expression of the latter is inhibited. According to these results, immunofluorescence experiments in Che-1-depleted cells showed an extensive accumulation of R-loops in the nucleoli, the main site of active transcription and where Che-1 is mostly found. In agreement with these results, RNA-seq experiments showed how the depletions of Che-1 or NEAT1 produce the activation of NF-kB and IFN pathways through the DNA-sensing receptor cyclic GMP– AMP synthase (cGAS) (53, 54), reinforcing the notion that these two molecules are involved in the resolution of RNA:DNA hybrids.

Recently, Che-1 has been shown to regulate chromatin accessibility(7). Further studies will define its role in the control of R-loops depends on this function or if these are two distinct functions of the protein.

It has been demonstrated that expression of both NEAT1 and Che-1 increases in response to several types of stress. The higher levels of NEAT1 are deemed necessary for a greater production of paraspeckles to sequester protein and RNA molecules and allow the cell to be able to respond adequately to the insult received. However, since most cellular stresses, as indeed viral infections, lead to an accumulation of RNA:DNA hybrids, it is also possible that Che-1 together with the other molecules of the paraspeckle, plays an active role on these hybrids, thus containing the generation of DNA damage and even cell death.

We further report increased levels of NEAT1 and paraspeckles genes during MM progression, and these data were further confirmed by clinical data available in CoMMpass data set. Moreover, the expression of NEAT1, Che-1, and other paraspeckles genes were found associated with reduced overall survival of MM patients. Remarkably, these findings concur with our finding that MM patients exhibit elevated levels of RNA:DNA hybrids compared to healthy controls, and this phenomenon appears to be related to disease progression, given that activation of IFN pathway is significantly found in ISS 3 and is markedly associated with a poor prognosis. It has recently been shown that cellular stress is capable of producing high levels of RNA:DNA hybrids(61). Therefore, the presence of a high quantity of these structures in MM patients could be produced by the constitutive UPR that distinguishes this pathology. In support of this hypothesis, our results from UPR modulation in MM cells confirmed a direct correlation between this type of stress, and the levels of RNA:DNA hybrids along with IFN response. Furthermore, it has recently been shown that proteasome inhibition induced by some drugs commonly used in the treatment of MM produces a broad-scale interference with spliceosome function (66), indicating a possible alternative molecular mechanism for the generation of RNA:DNA hybrids.

Our results show a direct correlation between MM progression and IFN activation. Sterile trigger of inflammation has been described to play an important role in the formation and maintenance of multipotent hematopoietic stem and progenitor cells (HSPCs) (67–69). It is very interesting to underline how a very recent study has identified the imbalance of R-loops as the cause of the activation of sterile inflammation in HSPCs, identifying an important link between R-loops, inflammation, and the development of the hematopoietic system (70). In addition, they found a strong correlation between inflammatory signaling and bone marrow blasts (70). It is therefore possible to hypothesize a model in which MM cells might take proliferative advantage from a high UPR, producing RNA:DNA hybrids, and triggering an inflammatory cascade (Figure 8). In this context, the elevated Che-1 and NEAT1 expression levels observed in MM, could help to keep the levels of RNA:DNA hybrids under constant control, preventing extensive genomic damage and cell death (Figure 8). Of course, further additional experiments are needed to understand the real relevance of these mechanisms in MM, but our results identify a new important role of Che-1 and NEAT1 in this pathology and reinforce the notion that these molecules can be considered important therapeutic targets.

**Figure 8.**
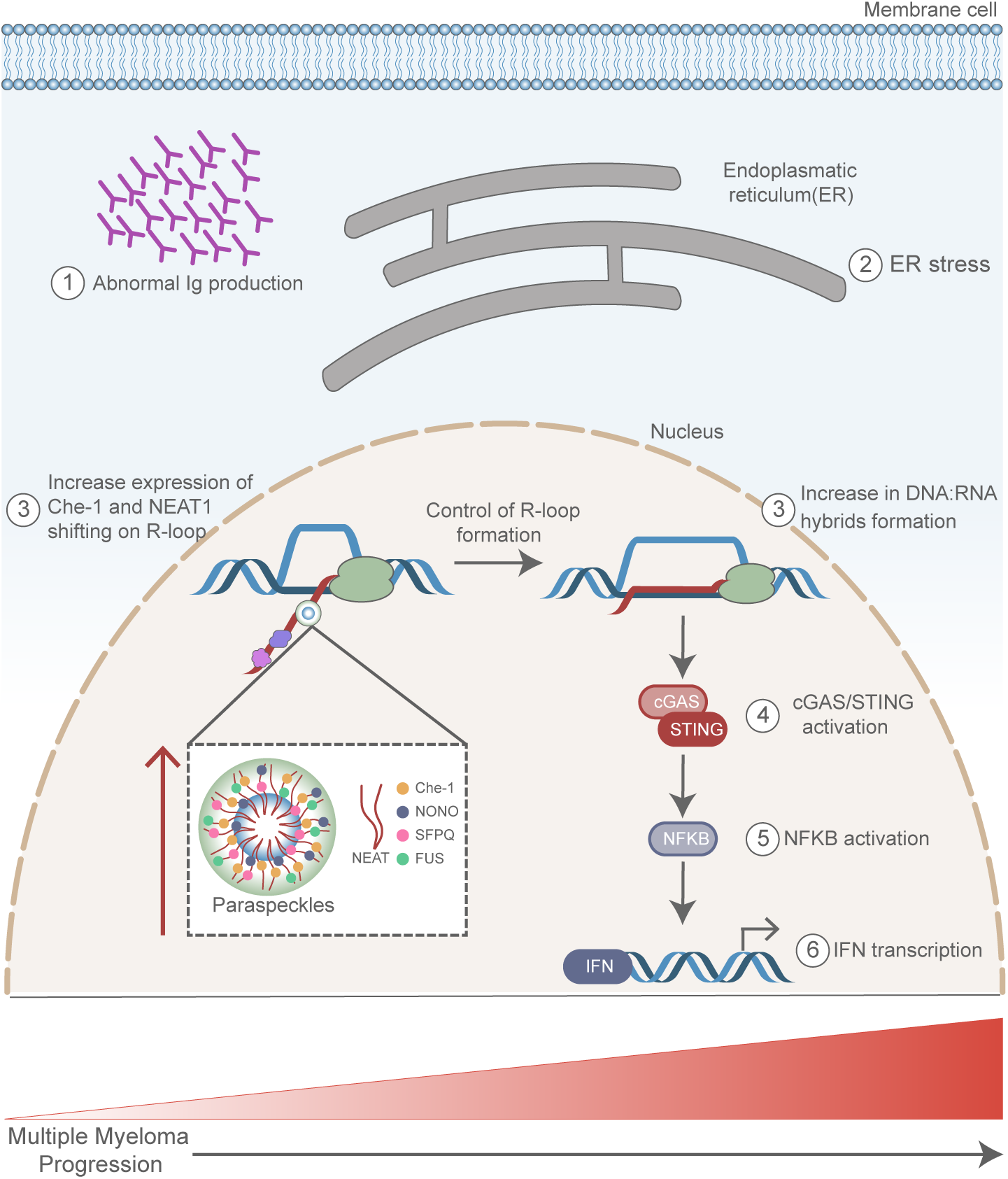
Model explaining INF activation in MM cells. (1) Abnormal production of immunoglobulin (Ig) in MM cells induces Endoplasmatic Reticulum (ER) stress (2). In response to this stress an increase of RNA:DNA hybrids (3) is observed, together with upregulation of Che-1, NEAT1 and Paraspeckles (3). RNA: DNA hybrids induce cGAS/Sting pathway activation (4), NF-kB induction (5) with a concomitant Interferon (IFN) response (6), which contributes to MM progression.

## Data Availability

The mass spectrometry data (Raw data and MaxQuant output) have been deposited to the Proteome Xchange Consortium (http://www.ebi.ac.uk/pride) via the PRIDE partner repository with the dataset identifier PXD026659.

Reviewer’s access:

Username

**Password: 6LfiJ05e**

High Throughput Sequencing data (RNA-seq, ChIP-seq, ChIRP-seq, DRIP-seq) from this publication have been submitted to the National Cancer Center for Biotechnology Information (NCBI) Gene Expression Omnibus database (https://www.ncbi.nlm.nih.gov/geo) and assigned the identifier numbers GSE178868. Reviewer’s access: https://www.ncbi.nlm.nih.gov/geo/query/acc.cgi?acc=GSE178868 using this private token: **avadmkaynjenvyl**

All other data supporting the findings of this study are available from the corresponding authors on reasonable request.

## Acknowledgments

We want to acknowledge and thank all patients and their families for the support and for donating the research samples. We thank Stefano Scalera and Marcello Maugeri-Saccà for the technical assistance in supervising the survival analysis. These data were generated as part of the Multiple Myeloma Research Foundation Personalized Medicine Initiatives (https://research.themmrf.org and www.themmrf.org).

## Correspondence

Maurizio Fanciulli, SAFU Laboratory, Department of Research, Advanced Diagnostics, and Technological Innovation, Translational Research Area, IRCCS Regina Elena National Cancer Institute, Via E. Chianesi 53, 00144, Rome, Italy., Tel.: (+3906) 5266-2800

## References

1. Kumar, S.K., Rajkumar, V., Kyle, R.A., van Duin, M., Sonneveld, P., Mateos, M.V., Gay, F. and Anderson, K.C. (2017) Multiple myeloma. Nat Rev Dis Primers, 3, 17046.

2. Manier, S., Salem, K., Glavey, S.V., Roccaro, A.M. and Ghobrial, I.M. (2016) Genomic Aberrations in Multiple Myeloma. Cancer Treat Res, 169, 23–34.

3. Manier, S., Salem, K.Z., Park, J., Landau, D.A., Getz, G. and Ghobrial, I.M. (2017) Genomic complexity of multiple myeloma and its clinical implications. Nat Rev Clin Oncol, 14, 100–113.

4. Neuse, C.J., Lomas, O.C., Schliemann, C., Shen, Y.J., Manier, S., Bustoros, M. and Ghobrial, I.M. (2020) Genome instability in multiple myeloma. Leukemia, 34, 2887–2897.

5. Cottini, F., Hideshima, T., Suzuki, R., Tai, Y.T., Bianchini, G., Richardson, P.G., Anderson, K.C. and Tonon, G. (2015) Synthetic Lethal Approaches Exploiting DNA Damage in Aggressive Myeloma. Cancer Discov, 5, 972–987.

6. Desantis, A., Bruno, T., Catena, V., De Nicola, F., Goeman, F., Iezzi, S., Sorino, C., Ponzoni, M., Bossi, G., Federico, V. et al. (2015) Che-1-induced inhibition of mTOR pathway enables stress-induced autophagy. EMBO J, 34, 1214–1230.

7. Bruno, T., De Nicola, F., Corleone, G., Catena, V., Goeman, F., Pallocca, M., Sorino, C., Bossi, G., Amadio, B., Cigliana, G. et al. (2020) Che-1/AATF-induced transcriptionally active chromatin promotes cell proliferation in multiple myeloma. Blood Adv, 4, 5616–5630.

8. Fanciulli, M., Bruno, T., Di Padova, M., De Angelis, R., Iezzi, S., Iacobini, C., Floridi, A. and Passananti, C. (2000) Identification of a novel partner of RNA polymerase II subunit 11, Che- 1, which interacts with and affects the growth suppression function of Rb. FASEB J, 14, 904–912.

9. Iezzi, S. and Fanciulli, M. (2015) Discovering Che-1/AATF: a new attractive target for cancer therapy. Front Genet, 6, 141.

10. Welcker, D., Jain, M., Khurshid, S., Jokic, M., Hohne, M., Schmitt, A., Frommolt, P., Niessen, C.M., Spiro, J., Persigehl, T. et al. (2018) AATF suppresses apoptosis, promotes proliferation and is critical for Kras-driven lung cancer. Oncogene, 37, 1503–1518.

11. Folgiero, V., Sorino, C., Pallocca, M., De Nicola, F., Goeman, F., Bertaina, V., Strocchio, L., Romania, P., Pitisci, A., Iezzi, S. et al. (2018) Che-1 is targeted by c-Myc to sustain proliferation in pre-B-cell acute lymphoblastic leukemia. EMBO Rep, 19.

12. Jing, P., Zou, J., Weng, K. and Peng, P. (2018) The PI3K/AKT axis modulates AATF activity in Wilms’ tumor cells. FEBS Open Bio, 8, 1615–1623.

13. Kumar, D.P., Santhekadur, P.K., Seneshaw, M., Mirshahi, F., Uram-Tuculescu, C. and Sanyal, A.J. (2019) A Regulatory Role of Apoptosis Antagonizing Transcription Factor in the Pathogenesis of Nonalcoholic Fatty Liver Disease and Hepatocellular Carcinoma. Hepatology, 69, 1520–1534.

14. Fox, A.H. and Lamond, A.I. (2010) Paraspeckles. Cold Spring Harb Perspect Biol, 2, a000687.

15. Fox, A.H., Nakagawa, S., Hirose, T. and Bond, C.S. (2018) Paraspeckles: Where Long Noncoding RNA Meets Phase Separation. Trends Biochem Sci, 43, 124–135.

16. Clemson, C.M., Hutchinson, J.N., Sara, S.A., Ensminger, A.W., Fox, A.H., Chess, A. and Lawrence, J.B. (2009) An architectural role for a nuclear noncoding RNA: NEAT1 RNA is essential for the structure of paraspeckles. Mol Cell, 33, 717–726.

17. Yamazaki, T. and Hirose, T. (2015) The building process of the functional paraspeckle with long non-coding RNAs. Front Biosci (Elite Ed), 7, 1–41.

18. West, J.A., Davis, C.P., Sunwoo, H., Simon, M.D., Sadreyev, R.I., Wang, P.I., Tolstorukov, M.Y. and Kingston, R.E. (2014) The long noncoding RNAs NEAT1 and MALAT1 bind active chromatin sites. Mol Cell, 55, 791–802.

19. Garcia-Muse, T. and Aguilera, A. (2019) R Loops: From Physiological to Pathological Roles. Cell, 179, 604–618.

20. Santos-Pereira, J.M. and Aguilera, A. (2015) R loops: new modulators of genome dynamics and function. Nat Rev Genet, 16, 583–597.

21. Belotserkovskii, B.P., Tornaletti, S., D’Souza, A.D. and Hanawalt, P.C. (2018) R-loop generation during transcription: Formation, processing and cellular outcomes. DNA Repair (Amst), 71, 69–81.

22. Groh, M. and Gromak, N. (2014) Out of balance: R-loops in human disease. PLoS Genet, 10, e1004630.

23. Bruno, T., Desantis, A., Bossi, G., Di Agostino, S., Sorino, C., De Nicola, F., Iezzi, S., Franchitto, A., Benassi, B., Galanti, S. et al. (2010) Che-1 promotes tumor cell survival by sustaining mutant p53 transcription and inhibiting DNA damage response activation. Cancer Cell, 18, 122–134.

24. Livak, K.J. and Schmittgen, T.D. (2001) Analysis of relative gene expression data using real-time quantitative PCR and the 2(-Delta Delta C(T)) Method. Methods, 25, 402–408.

25. Sorino, C., Catena, V., Bruno, T., De Nicola, F., Scalera, S., Bossi, G., Fabretti, F., Mano, M., De Smaele, E., Fanciulli, M. et al. (2020) Che-1/AATF binds to RNA polymerase I machinery and sustains ribosomal RNA gene transcription. Nucleic Acids Res, 48, 5891–5906.

26. Chu, C., Quinn, J. and Chang, H.Y. (2012) Chromatin isolation by RNA purification (ChIRP). J Vis Exp.

27. Ginno, P.A., Lott, P.L., Christensen, H.C., Korf, I. and Chedin, F. (2012) R-loop formation is a distinctive characteristic of unmethylated human CpG island promoters. Mol Cell, 45, 814–825.

28. Ewels, P., Magnusson, M., Lundin, S. and Kaller, M. (2016) MultiQC: summarize analysis results for multiple tools and samples in a single report. Bioinformatics, 32, 3047–3048.

29. Langmead, B. and Salzberg, S.L. (2012) Fast gapped-read alignment with Bowtie 2. Nat Methods, 9, 357–359.

30. Li, H., Handsaker, B., Wysoker, A., Fennell, T., Ruan, J., Homer, N., Marth, G., Abecasis, G., Durbin, R. and Genome Project Data Processing, S. (2009) The Sequence Alignment/Map format and SAMtools. Bioinformatics, 25, 2078–2079.

31. Kent, W.J., Zweig, A.S., Barber, G., Hinrichs, A.S. and Karolchik, D. (2010) BigWig and BigBed: enabling browsing of large distributed datasets. Bioinformatics, 26, 2204–2207.

32. Heinz, S., Benner, C., Spann, N., Bertolino, E., Lin, Y.C., Laslo, P., Cheng, J.X., Murre, C., Singh, H. and Glass, C.K. (2010) Simple combinations of lineage-determining transcription factors prime cis-regulatory elements required for macrophage and B cell identities. Mol Cell, 38, 576–589.

33. Yates, A.D., Achuthan, P., Akanni, W., Allen, J., Allen, J., Alvarez-Jarreta, J., Amode, M.R., Armean, I.M., Azov, A.G., Bennett, R. et al. (2020) Ensembl 2020. Nucleic Acids Res, 48, D682–D688.

34. Ramirez, F., Dundar, F., Diehl, S., Gruning, B.A. and Manke, T. (2014) deepTools: a flexible platform for exploring deep-sequencing data. Nucleic Acids Res, 42, W187–191.

35. Bray, N.L., Pimentel, H., Melsted, P. and Pachter, L. (2016) Near-optimal probabilistic RNA-seq quantification. Nat Biotechnol, 34, 525–527.

36. Pimentel, H., Bray, N.L., Puente, S., Melsted, P. and Pachter, L. (2017) Differential analysis of RNA-seq incorporating quantification uncertainty. Nat Methods, 14, 687–690.

37. Durinck, S., Spellman, P.T., Birney, E. and Huber, W. (2009) Mapping identifiers for the integration of genomic datasets with the R/Bioconductor package biomaRt. Nat Protoc, 4, 1184–1191.

38. Schubert, M., Klinger, B., Klunemann, M., Sieber, A., Uhlitz, F., Sauer, S., Garnett, M.J., Bluthgen, N. and Saez-Rodriguez, J. (2018) Perturbation-response genes reveal signaling footprints in cancer gene expression. Nat Commun, 9, 20.

39. Robinson, M.D., McCarthy, D.J. and Smyth, G.K. (2010) edgeR: a Bioconductor package for differential expression analysis of digital gene expression data. Bioinformatics, 26, 139–140.

40. Barwick, B.G., Neri, P., Bahlis, N.J., Nooka, A.K., Dhodapkar, M.V., Jaye, D.L., Hofmeister, C.C., Kaufman, J.L., Gupta, V.A., Auclair, D. et al. (2019) Multiple myeloma immunoglobulin lambda translocations portend poor prognosis. Nat Commun, 10, 1911.

41. Bolli, N., Avet-Loiseau, H., Wedge, D.C., Van Loo, P., Alexandrov, L.B., Martincorena, I., Dawson, K.J., Iorio, F., Nik-Zainal, S., Bignell, G.R. et al. (2014) Heterogeneity of genomic evolution and mutational profiles in multiple myeloma. Nat Commun, 5, 2997.

42. Kaiser, R.W.J., Ignarski, M., Van Nostrand, E.L., Frese, C.K., Jain, M., Cukoski, S., Heinen, H., Schaechter, M., Seufert, L., Bunte, K. et al. (2019) A protein-RNA interaction atlas of the ribosome biogenesis factor AATF. Sci Rep, 9, 11071.

43. Liu, X., Cai, S., Zhang, C., Liu, Z., Luo, J., Xing, B. and Du, X. (2018) Deacetylation of NAT10 by Sirt1 promotes the transition from rRNA biogenesis to autophagy upon energy stress. Nucleic Acids Res, 46, 9601–9616.

44. Bammert, L., Jonas, S., Ungricht, R. and Kutay, U. (2016) Human AATF/Che-1 forms a nucleolar protein complex with NGDN and NOL10 required for 40S ribosomal subunit synthesis. Nucleic Acids Res, 44, 9803–9820.

45. Van Nostrand, E.L., Freese, P., Pratt, G.A., Wang, X., Wei, X., Xiao, R., Blue, S.M., Chen, J.Y., Cody, N.A.L., Dominguez, D. et al. (2020) A large-scale binding and functional map of human RNA-binding proteins. Nature, 583, 711–719.

46. Yamazaki, T., Souquere, S., Chujo, T., Kobelke, S., Chong, Y.S., Fox, A.H., Bond, C.S., Nakagawa, S., Pierron, G. and Hirose, T. (2018) Functional Domains of NEAT1 Architectural lncRNA Induce Paraspeckle Assembly through Phase Separation. Mol Cell, 70, 1038–1053 e1037.

47. Hirose, T., Virnicchi, G., Tanigawa, A., Naganuma, T., Li, R., Kimura, H., Yokoi, T., Nakagawa, S., Benard, M., Fox, A.H. et al. (2014) NEAT1 long noncoding RNA regulates transcription via protein sequestration within subnuclear bodies. Mol Biol Cell, 25, 169–183.

48. Cristini, A., Groh, M., Kristiansen, M.S. and Gromak, N. (2018) RNA/DNA Hybrid Interactome Identifies DXH9 as a Molecular Player in Transcriptional Termination and R-Loop-Associated DNA Damage. Cell Rep, 23, 1891–1905.

49. Boguslawski, S.J., Smith, D.E., Michalak, M.A., Mickelson, K.E., Yehle, C.O., Patterson, W.L. and Carrico, R.J. (1986) Characterization of monoclonal antibody to DNA.RNA and its application to immunodetection of hybrids. J Immunol Methods, 89, 123–130.

50. Sanz, L.A., Hartono, S.R., Lim, Y.W., Steyaert, S., Rajpurkar, A., Ginno, P.A., Xu, X. and Chedin, F. (2016) Prevalent, Dynamic, and Conserved R-Loop Structures Associate with Specific Epigenomic Signatures in Mammals. Mol Cell, 63, 167–178.

51. Shen, Y.J., Le Bert, N., Chitre, A.A., Koo, C.X., Nga, X.H., Ho, S.S., Khatoo, M., Tan, N.Y., Ishii, K.J. and Gasser, S. (2015) Genome-derived cytosolic DNA mediates type I interferon-dependent rejection of B cell lymphoma cells. Cell Rep, 11, 460–473.

52. Lim, Y.W., Sanz, L.A., Xu, X., Hartono, S.R. and Chedin, F. (2015) Genome-wide DNA hypomethylation and RNA:DNA hybrid accumulation in Aicardi-Goutieres syndrome. Elife, 4.

53. Hopfner, K.P. and Hornung, V. (2020) Molecular mechanisms and cellular functions of cGAS-STING signalling. Nat Rev Mol Cell Biol, 21, 501–521.

54. Motwani, M., Pesiridis, S. and Fitzgerald, K.A. (2019) DNA sensing by the cGAS-STING pathway in health and disease. Nat Rev Genet, 20, 657–674.

55. Yoneyama, M., Kikuchi, M., Natsukawa, T., Shinobu, N., Imaizumi, T., Miyagishi, M., Taira, K., Akira, S. and Fujita, T. (2004) The RNA helicase RIG-I has an essential function in double-stranded RNA-induced innate antiviral responses. Nat Immunol, 5, 730–737.

56. Nolan, G.P., Ghosh, S., Liou, H.C., Tempst, P. and Baltimore, D. (1991) DNA binding and I kappa B inhibition of the cloned p65 subunit of NF-kappa B, a rel-related polypeptide. Cell, 64, 961–969.

57. Agnelli, L., Mosca, L., Fabris, S., Lionetti, M., Andronache, A., Kwee, I., Todoerti, K., Verdelli, D., Battaglia, C., Bertoni, F. et al. (2009) A SNP microarray and FISH-based procedure to detect allelic imbalances in multiple myeloma: an integrated genomics approach reveals a wide gene dosage effect. Genes Chromosomes Cancer, 48, 603–614.

58. Taiana, E., Favasuli, V., Ronchetti, D., Todoerti, K., Pelizzoni, F., Manzoni, M., Barbieri, M., Fabris, S., Silvestris, I., Gallo Cantafio, M.E. et al. (2020) Long non-coding RNA NEAT1 targeting impairs the DNA repair machinery and triggers anti-tumor activity in multiple myeloma. Leukemia, 34, 234–244.

59. Taiana, E., Ronchetti, D., Favasuli, V., Todoerti, K., Manzoni, M., Amodio, N., Tassone, P., Agnelli, L. and Neri, A. (2019) Long non-coding RNA NEAT1 shows high expression unrelated to molecular features and clinical outcome in multiple myeloma. Haematologica, 104, e72–e76.

60. Hanamura, I., Huang, Y., Zhan, F., Barlogie, B. and Shaughnessy, J. (2006) Prognostic value of cyclin D2 mRNA expression in newly diagnosed multiple myeloma treated with high-dose chemotherapy and tandem autologous stem cell transplantations. Leukemia, 20, 1288–1290.

61. Choo, J., Schlosser, D., Manzini, V., Magerhans, A. and Dobbelstein, M. (2020) The integrated stress response induces R-loops and hinders replication fork progression. Cell Death Dis, 11, 538.

62. Nikesitch, N., Lee, J.M., Ling, S. and Roberts, T.L. (2018) Endoplasmic reticulum stress in the development of multiple myeloma and drug resistance. Clin Transl Immunology, 7, e1007.

63. Sidrauski, C., Acosta-Alvear, D., Khoutorsky, A., Vedantham, P., Hearn, B.R., Li, H., Gamache, K., Gallagher, C.M., Ang, K.K., Wilson, C. et al. (2013) Pharmacological brake-release of mRNA translation enhances cognitive memory. Elife, 2, e00498.

64. Tsai, J.C., Miller-Vedam, L.E., Anand, A.A., Jaishankar, P., Nguyen, H.C., Renslo, A.R., Frost, A. and Walter, P. (2018) Structure of the nucleotide exchange factor eIF2B reveals mechanism of memory-enhancing molecule. Science, 359.

65. Zyryanova, A.F., Weis, F., Faille, A., Alard, A.A., Crespillo-Casado, A., Sekine, Y., Harding, H.P., Allen, F., Parts, L., Fromont, C. et al. (2018) Binding of ISRIB reveals a regulatory site in the nucleotide exchange factor eIF2B. Science, 359, 1533–1536.

66. Huang, H.H., Ferguson, I.D., Thornton, A.M., Bastola, P., Lam, C., Lin, Y.T., Choudhry, P., Mariano, M.C., Marcoulis, M.D., Teo, C.F. et al. (2020) Proteasome inhibitor-induced modulation reveals the spliceosome as a specific therapeutic vulnerability in multiple myeloma. Nat Commun, 11, 1931.

67. Espin-Palazon, R., Weijts, B., Mulero, V. and Traver, D. (2018) Proinflammatory Signals as Fuel for the Fire of Hematopoietic Stem Cell Emergence. Trends Cell Biol, 28, 58–66.

68. Pietras, E.M. (2017) Inflammation: a key regulator of hematopoietic stem cell fate in health and disease. Blood, 130, 1693–1698.

69. Sawamiphak, S., Kontarakis, Z. and Stainier, D.Y. (2014) Interferon gamma signaling positively regulates hematopoietic stem cell emergence. Dev Cell, 31, 640–653.

70. Weinreb, J.T., Gupta, V., Sharvit, E., Weil, R. and Bowman, T.V. (2021) Ddx41 inhibition of DNA damage signaling permits erythroid progenitor expansion in zebrafish. Haematologica.

