## supplemental figures for "AATF/Che-1, a new component of paraspeckles, controls R-loops formation and Interferon activation in Multiple Myeloma"

**Contents:**

- Legend of Supplementary Figures
- Supplementary References
- Supplementary Table 2 Antibodies index
- Supplementary Table 3 Primers index
- Supplementary Table 4
- Supplementary Figures

### Supplementary Figures Legends

**Supplementary Figure S1, related to Figure 1. Che-1 interacts with paraspeckle components.**

**A:** Scatter Plot of mass spectrometry assay in Hela cells(1). Dots depict protein intensity (IBAQ) and relative significance obtained from mass spectrometry assay. X-axis: Difference of Che-1 intensity versus IP control; on the y-axis are the corresponding −log10 P-values. Significant Che-1 interacting proteins relevant in this study are highlighted in blue. **B:** Heatmap showing the eCLIP binding sites of 120 RBPs at NEAT1 domains(2). The binding is defined by intersecting the eCLIP profiles of each RBP with the NEAT1 domains and assigning 1 in case of overlaps are found or 0 otherwise.

**Supplementary Figure S2, related to Figure 2. Che-1 and NEAT1 colocalize on the DNA.**

**A:** Barplot indicating the number of enriched sites in ChIRP-seq NEAT1 (yellow) and ChIP-seq (red) experiments corresponding respectively to 19.137 and 41.061 sites. **B:** Venn diagram showing shared and unique peaks in 3 pairwise comparisons: H3K27ac vs. co-localizing sites (right), H3K27ac vs. only AATF/Che1 sites (middle) and H3K27ac vs. only NEAT1 sites (left).

**Supplementary Figure S3, related to Figure 3. NEAT1 is required for recruiting Che-1 onto the DNA.**

**A:** Representative images of RPMI8226 MM cells transfected with GapmeR NEAT1 or GapmeR Control and immunostained with specific Che-1 antibody. One of two experiments is shown. Scale bar 10 μm. **B:** Cell number–normalized total RNA quantification of Kms27 MM cells depleted or not for NEAT1 GapmeR oligonucleotide. Data represent the mean ± SD. Statistical significance is indicated by asterisks as follows: *$P$ 0.046. Statistical analysis was performed using two-sided t-tests. **C:** Measurement of Kms27 MM cell viability after transfection with NEAT1 or control GapmeR oligolucleotides. Error bars represent the SD of triplicate experiments. Statistical significance is indicated by asterisks as follows: *$P$ 0.0003. Statistical analysis was performed using two-sided t-tests. **D:** RT-qPCR analysis for the indicated genes from Kms27 cells transfected with NEAT1 or control GapmeR oligonucleotides. Results from three biological replicates are shown. Statistical significance is indicated by asterisks as follows: *$P$< 0.049. Statistical analysis was performed using two-sided t-tests.

**Supplementary Figure S4, related to Figure 4. Che-1 and NEAT1 suppress RNA: DNA hybrids accumulation.**

**A:** RIP assay performed in RPMI8226 and Kms27MM cells using an anti-Che-1 specific antibody. IgG were used as negative control. NEAT1 RNA abundance was evaluated by RT–qPCR. Results from at least three biological replicates for each line are shown. Statistical significance is indicated by asterisks as follows: *$P$< 0.018. Statistical analysis was performed using two-sided t-tests. **B**: Immunostaining of Kms27 MM cells with S9.6 antibody (green) and Hoechst (blue), transfected with LNA GapmeR Control or GapmeR NEAT1 oligonucleotides and treated or not with RNase H. Scale bar 10 μM. **C:** DRIP-qPCR analysis of indicated genes in Kms27 MM cells treated or not with RNase H. Results from three biological replicates are shown. Statistical significance is indicated by asterisks as follows: *$P$< 0.01. Statistical analysis was performed using two-sided t-tests.

**Supplementary Figure S5, related to Figure 5. Che-1 controls Interferon activation in MM cells.**

**A:** Differential analysis of GapmeR Control vs. GapmeR NEAT1 transcriptome in Kms27 MM cells. Volcano plot shows 174 significantly upregulated (red) and 41 downregulated genes (light blue). x-axis reports the base-2 logarithm of fold change which is approximated to the B value coming from the statistical Wald test. y-axis reports the base-10 logarithm of Q value of significant genes. **B:** Gene Set Enrichment Analysis (GSEA) of upregulated significant differential genes with their relative base-10 logarithm of Q value. The analysis was based on hallmark gene sets collection. **c:** Heatmap depicting pathway activity scores for each transcriptome sample according to PROGENy. The activity score is shown as shade ranging from blue to red which indicates the level of pathway activation. Blu= non-active; Red=active. **D, E, and F:** RT-qPCR of the indicated genes was performed after transient transfection of Kms27 MM cells with GapmeR Control or NEAT1 GapmeR oligonucleotides. Error bars represent the standard error of three different experiments. Statistical significance is indicated by asterisks as follows: $*P< 0.02$, $**P< 0.05$. Statistical analysis was performed using two-sided t-tests.

**Supplementary Figure S6, related to Figure 6. MM patients exhibit high levels of RNA: DNA hybrids.**

**A:** Data from Agnelli et al.(3) provided by Oncomine database and reanalyzed for the expression of NEAT1, SFPQ and NONO, in normal bone marrow, MGUS, MM and plasma cell leukemia ($n$ = 158). The associate $P$-value is shown above Box and whisker plot shows the upper and lower quartiles (25-75%) with a line at the median, whiskers extend from $10^{th}$ to the $90^{th}$ percentile, and dots correspond to the minimal and maximal values. **B:** Boxplot showing the relative enrichment of paraspeckles genes (right) and NEAT1 (left), derived from the transcriptomes of the selected CoMMpass patient cohort (N=687), by the relative ISS score. The enrichment values reported on y-axis correspond to raw reads counts normalized with weighted trimmed mean of M-values (TMM). **c:** Kaplan-Meier survival curve for NEAT1, SFPQ and NONO expression in the Hanamura et al.(4) cohort of MM patients (n = 542).

**Supplementary Figure S7, related to Figure 7. UPR induced R-loops in MM cells.**

**A:** NEAT1 levels in Kms27 and U266 MM cells were evaluated by RT-qPCR analysis. Data are presented as mean ± SD of three independent experiments. Statistical significance is indicated by asterisks as follows: $*P$ 7.8E-08. Statistical analysis was performed using two-sided t-tests. **B**: S9.6 antibody was used for dot blot analysis to evaluated RNA:DNA hybrids formation in both MM cells Kms27 and U266 by serial dilutions of genomic DNA starting at 1.5 micrograms. Methylene blue normalization of S9.6 signal image was acquired by using Alliance Mini HD6 system by UVITEC Ltd,. One out of two experiments are shown. **C:** RT-qPCR was performed to assessed NEAT1 level in U266 MM cells treated or not for THAP 0,5μM for 16 hours. Error bars represented the results from three independent experiments. Statistical significance is indicated by asterisks as follows: *$P$ 1,55E-05. Statistical analysis was performed using two-sided t-tests. **D:** U266 MM cells treated or not for THAP 0,5μM for 16 hours was used for dot blot analysis of RNA:DNA hybrids formation as in **B**. Methylene blue normalization of S9.6 signal image was acquired by using Alliance Mini HD6 system by UVITEC Ltd,. One out of two experiments are shown. **E:** RT-qPCR analysis of NEAT1 levels in Kms27 MM cells treated or not with ISRIB 1μM for 6 hours. Results from three biological replicates are shown. Statistical significance is indicated by asterisks as follows: *$P$ 3,24E-05. Statistical analysis was performed using two-sided t-tests. **F:** Kms27 MM cells exposed or not to ISRIB 1μM for 6 hours were used for dot blot analysis of RNA:DNA hybrids formation as in **B**. Methylene blue normalization of S9.6 signal image was acquired by using Alliance Mini HD6 system by UVITEC Ltd,. One out of two experiments are shown.

**References**

1. Kaiser, R.W.J., Ignarski, M., Van Nostrand, E.L., Frese, C.K., Jain, M., Cukoski, S., Heinen, H., Schaechter, M., Seufert, L., Bunte, K. *et al.* (2019) A protein-RNA interaction atlas of the ribosome biogenesis factor AATF. *Sci Rep*, **9**, 11071.
2. Van Nostrand, E.L., Freese, P., Pratt, G.A., Wang, X., Wei, X., Xiao, R., Blue, S.M., Chen, J.Y., Cody, N.A.L., Dominguez, D. *et al.* (2020) A large-scale binding and functional map of human RNA-binding proteins. *Nature*, **583**, 711-719.
3. Agnelli, L., Mosca, L., Fabris, S., Lionetti, M., Andronache, A., Kwee, I., Todoerti, K., Verdelli, D., Battaglia, C., Bertoni, F. *et al.* (2009) A SNP microarray and FISH-based procedure to detect allelic imbalances in multiple myeloma: an integrated genomics approach reveals a wide gene dosage effect. *Genes Chromosomes Cancer*, **48**, 603-614.
4. Hanamura, I., Huang, Y., Zhan, F., Barlogie, B. and Shaughnessy, J. (2006) Prognostic value of cyclin D2 mRNA expression in newly diagnosed multiple myeloma treated with high-dose chemotherapy and tandem autologous stem cell transplantations. *Leukemia*, **20**, 1288-1290.

### Supplementary Table S2: Antibodies index

| Antibodies | SOURCE | IDENTIFIERS |
|---|---|---|
| Rabbit Polyclonal anti-Che-1 | Millipore | Cat# ABC953 |
| Rabbit Polyclonal anti-Histone H3 | Abcam | Cat# ab1791 |
| Rabbit Polyclonal anti-Acetyl Histone H3 | Millipore | Cat# 06-599 |
| Mouse Monoclonal anti-β-actin | Sigma Aldrich | Cat# A5441 |
| Spike-in Antibody | Active Motif | Cat# 61686 |
| Rabbit Polyclonal anti-SFPQ | Bethyl | Cat#A301-322A |
| Rabbit Polyclonal anti-NONO | Bethyl | Cat#A300-587A |
| Mouse Monoclonal anti SFPQ | ThermoFisher | Cat#MA1-25325 |
| Mouse Monoclonal anti-NONO (78-1 C) | ThermoFisher | Cat#MA3-2024 |
| Rabbit Polyclonal anti-cGAS (E5V3W) | Cell Signaling | Cat#79978 |
| Rabbit Polyclonal anti-RIG1 (D33H10) | Cell Signaling | Cat#4200 |
| Rabbit Polyclonal anti-ATF4 (D4B8) | Cell Signaling | Cat#11815 |
| Rabbit Polyclonal anti-PDI (C81H6) | Cell Signaling | Cat#3501 |
| Rabbit Polyclonal anti-PERK (D11A8) | Cell Signaling | Cat#5683 |
| Rabbit Polyclonal anti-IRE1a (14C10) | Cell Signaling | Cat#3294 |
| Rabbit Polyclonal anti-CHOP (D46F1) | Cell Signaling | Cat#5554 |
| Rabbit Polyclonal anti-GFP (D5.1) | Cell Signaling | Cat#2956 |
| Mouse Monoclonal anti-S9.6 | Kerafast | Cat#ENH001 |
| Rabbit Polyclonal anti-STING (D2P2F) | Cell Signaling | Cat#13647 |

**Supplemental Table S3 – Primers Index**

| **RT-qPCR mRNA primers** | | |
|---|---|---|
| **Human Primers** | **Fwd/Rvs** | **Sequence 5'- 3'** |
| Actin | Fwd | GACAGGATGCAGAAGGAGATTACT |
| | Rvs | TGATCCACATCTGCTGGAAGGT |
| OASL | Fwd | CGGGTGCTGAAGGTAGTCAA |
| | Rvs | AAACAGCTCAGAAACGCCAC |
| ISG15 | Fwd | GCGAACTCATCTTTGCCAGTA |
| | Rvs | CCAGCATCTTCACCGTCAG |
| DDX60 | Fwd | TGGATGCGTTGAATTATAGACAG |
| | Rvs | GAAATATACATCTCCCATCAGGTCTT |
| NEAT1_1 | Fwd | TGGCTAGCTCAGGGCTTCAG |
| | Rvs | TCTCCTTGCCAAGCTTCCTTC |
| NEAT1_2 | Fwd | CTAGAGGCTCGCATTGTGTG |
| | Rvs | GCCCACACGAAACCTTACAT |
| IRF8 | Fwd | ACGCTGTGCTTTGAATAAGAGC |
| | Rvs | TCCTCAGGAACAATTCGGTAAAC |
| ICAM1 | Fwd | ATGCCCAGACATCTGTGTCC |
| | Rvs | GGGGTCTCTATGCCCAACAA |
| TNFAIP3 | Fwr | TCCTCAGGCTTTGTATTTGAGC |
| | Rvs | TGTGTATCGGTGCATGGTTTTA |
| IFN β | Fwd | CCTGTGGCAATTGAATGGGAGGC |
| | Rvs | CCAGGCACAGTGACTGTACTCCTT |
| IFN γ | Fwd | TTCGGTAACTGACTTGAATGTCCA |
| | Rvs | TTTCGCTTCCCTGTTTTAGCT-G |
| Mycoplasm | Fwd | ACTCCTACGGGAGGCAGCAGTA |
| | Rvs | TCGACCATCTGTCACTCTGTTAAC |
| Epstein-Barr virus (EBV) | Fwd | GGAACCTGGTCATCCTTTGC |
| | Rvs | ACGTGCATGGACCGGTTAAT |
| RPL13A | Fwd | AGGTGCCTTGCTCACAGAGT |
| | Rvs | GGTTGCATTGCCCTCATTAC |
| FBXL17 | Fwd | CACCCTCGGAATCCTGTCTA |
| | Rvs | GCCTGCATAGCTGTTTCCTC |
| TRIM33 | Fwd | ATGCCCAGCTTTCCCTAACT |
| | Rvs | GGAAAGTGGACTGCATGGTT |
| Negative Control | Fwd | GAACGTTCAGCCTCGTTCTC |
| | Rvs | GGAAGGTGGAAGGAAACACA |
| **ChIP-qPCR primers** | **Fwd/Rvs** | **Sequence** |
| Che-1 | Fwd | CGCGCGCATCGCAATCGCATC |
| | Rvs | CGTCACTGCGGGCGTTGCTAG |
| Drosophila Spike-in | Fwd | AGGTGCTCCTTCAGGCGATTC |
| | Rvs | TATCCGGTGTTCATCGTGTAGC |
| **Capture oligo CHART seq** | | |
| NEAT1 CO1 | | TTCCTTCTCGCACCCCCAGC/iSp18//3BioTEG |
| NEAT1 CO2 | | TGTCTGTCCCCTGAAGCCCTG/iSp18//3BioTEG |
| NEAT1 CO3 | | CTAGCCACTTCCTCCCCCACAA/iSp18//3BioTEG |
| NEAT1 sense CO | | GCTGGGGGTGCGAGAAGGAA/iSp18//3BioTEG |

**Supplemental Table S4 Distribution of variables in reference population.**

| Characteristic | N (%) |
|---|---|
| **Median Age** | 62(56-69) |
| **ISS** | |
| **I** | 226 |
| **II** | 228 |
| **III** | 178 |
| **Therapy** | |
| **K-based** | 43 |
| **Combo K/IMIDs-based** | 286 |
| **Combo V/IMIDs/K-based** | 21 |
| **Combo IMIDs/K-based** | 116 |
| **IMIDs-based** | 34 |
| **IFN** | |
| **high** | 341 |
| **low** | 291 |
| **ASCT** | |
| **Yes** | 313 |
| **No** | 271 |
| **Nonvaluable** | 48 |
| **ECOG** | |
| **0** | 150 |
| **1** | 220 |
| **≥2** | 74 |
| **Missing** | 188 |
| **t (11;14)** | |
| **Yes** | 112 |
| **No** | 454 |
| **Nonvaluable** | 66 |
| **t (4;14)** | |
| **Yes** | 68 |
| **No** | 498 |
| **Nonvaluable** | 66 |
| **t (14;16)** | |
| **Yes** | 23 |
| **No** | 543 |
| **Nonvaluable** | 66 |
| **t (14;20)** | |
| **Yes** | 7 |
| **No** | 559 |
| **Nonvaluable** | 66 |
| **FGFR3** | |
| **Yes** | 21 |
| **No** | 572 |
| **Missing** | 39 |
| **SP140** | |
| **Yes** | 577 |
| **No** | 16 |
| **Missing** | 39 |
| **DIS3** | |
| **Yes** | 65 |
| **No** | 528 |

| | |
|---|---|
| **Missing** | 39 |
| **CYLD**<br>**Yes**<br>**No**<br>**Missing** | <br>23<br>570<br>39 |
| **HUWE1**<br>**Yes**<br>**No**<br>**Missing** | <br>18<br>575<br>39 |
| **PRKD2**<br>**Yes**<br>**No**<br>**Missing** | <br>18<br>575<br>39 |
| **EGR1**<br>**Yes**<br>**No**<br>**Missing** | <br>21<br>571<br>39 |
| **MAX**<br>**Yes**<br>**No**<br>**Missing** | <br>18<br>575<br>39 |
| **TRAF3**<br>**Yes**<br>**No**<br>**Missing** | <br>45<br>548<br>39 |
| **KRAS**<br>**Yes**<br>**No**<br>**Missing** | <br>154<br>439<br>39 |
| **TP53**<br>**Yes**<br>**No**<br>**Missing** | <br>26<br>567<br>39 |
| **ATM**<br>**Yes**<br>**No**<br>**Missing** | <br>15<br>578<br>39 |
| **BRAF**<br>**Yes**<br>**No**<br>**Missing** | <br>48<br>545<br>39 |
| **DUSP2**<br>**Yes**<br>**No**<br>**Missing** | <br>30<br>563<br>39 |
| **FAT3**<br>**Yes**<br>**No**<br>**Missing** | <br>33<br>560<br>39 |
| **HIST1H1E**<br>**Yes**<br>**No**<br>**Missing** | <br>29<br>564<br>39 |
| **CKS1B**<br>**Yes** | |

| | |
|---|---|
| **No**<br>**Missing** | |
| **FAM46C**<br>**Yes**<br>**No**<br>**Missing** | <br>64<br>529<br>39 |
| **ACTG1**<br>**Yes**<br>**No**<br>**Missing** | <br>14<br>579<br>39 |
| **NRAS**<br>**Yes**<br>**No**<br>**Missing** | <br>133<br>460<br>39 |
| **LTB**<br>**Yes**<br>**No**<br>**Missing** | <br>19<br>574<br>39 |
| **IGLL5**<br>**Yes**<br>**No**<br>**Missing** | <br>96<br>497<br>39 |

### List of classification method for each analysed variable.

| Variable | Categories | Method |
|---|---|---|
| **IFN** | High/low | RNA-seq |
| **ISS** | I, II, III | Baseline albumin, 2microglobulin[22] |
| **Therapy** | V-based, K-based, combo K/IMIDs-based, combo V/IMIDs/K-based, combo IMIDs/K-based, IMIDs-based | Therapy classification |
| **Translocation** | Presence/absence | Seq-FISH |
| **nsSNV/INDEL in a custom 21 genes panel** | Presence/absence | Whole-exome sequencing |
| **ASCT** | Yes/No | Therapy classification |
| **ECOG** | 0, 1,≥ 2 | Baseline ECOG |
| **Gender** | F/M | Baseline age |

### Statistics measurements from univariate analysis for each variable

| Variable | **log-rank pvalue** | **wd pvalue** | **beta** | **HR (95% CI for HR)** |
|---|---|---|---|---|
| **Age** | 7.9e-06 | 7.4e-06 | 0.035 | 1 (1-1.1) |
| **Gender** | 0.0091 | 0.0097 | -0.45 | 0.64 (0.46-0.9) |
| **ASCT** | 1.5e-10 | 9.3e-10 | -1.1 | 0.35 (0.25-0.49) |
| **ISS** | 1.2e-09 | 3.4e-09 | 0.62 | 1.8 (1.5-2.3) |

| | | | | |
|---|---|---|---|---|
| **IFN** | 1.9e-05 | 2.7e-05 | -0.71 | 0.49 (0.36-0.69) |
| **ECOG** | 0.00039 | 0.00041 | 0.35 | 1.4 (1.2-1.7) |
| **Therapy**<br>K-based<br>K/IMIDs-based<br>V/IMIDs/K-based<br>IMIDs/K-based<br>IMIDs-based | <br>0.12643<br>1.03e-05<br>0.00716<br>3.51e-05<br>0.42191 | 2e-06 | <br>-0.7959<br>-0.7837<br>-1.5971<br>-1.7829<br>-0.2490 | <br>0.4512<br>0.4567<br>0.2025<br>0.1682<br>0.7796 |
| **CCND1 translocation** | 0.45 | 0.45 | -0.17 | 0.85 (0.54-1.3) |
| **WHSC1 translocation** | 0.035 | 0.036 | 0.48 | 1.6 (1-2.5) |
| **MAF translocation** | 0.2 | 0.21 | 0.46 | 1.6 (0.78-3.2) |
| **MAFB translocation** | 0.18 | 0.19 | 0.76 | 2.1 (0.68-6.7) |
| **MYC translocation** | 0.11 | 0.11 | 0.34 | 1.4 (0.92-2.1) |
| **CCND2 translocation** | 0.8 | 0.8 | -0.25 | 0.78 (0.11-5.6) |
| **MAFA translocation** | 0.33 | 0.35 | -0.94 | 0.39 (0.055-2.8) |
| **FGFR3** | 0.062 | 0.067 | 0.67 | 2 (0.95-4) |
| **SP140** | 0.82 | 0.82 | 0.12 | 1.1 (0.42-3) |
| **DIS3** | 0.12 | 0.12 | 0.39 | 1.5 (0.91-2.4) |
| **CYLD** | 0.2 | 0.21 | -0.73 | 0.48 (0.15-1.5) |
| **HUWE1** | 0.3 | 0.31 | -0.73 | 0.48 (0.12-2) |
| **PRKD2** | 0.46 | 0.46 | 0.33 | 1.4 (0.57-3.4) |
| **EGR1** | 0.79 | 0.79 | -0.13 | 0.88 (0.32-2.4) |
| **MAX** | 0.87 | 0.87 | 0.077 | 1.1 (0.44-2.6) |
| **TRAF3** | 0.15 | 0.16 | -0.55 | 0.58 (0.27-1.2) |
| **KRAS** | 0.026 | 0.027 | 0.39 | 1.5 (1-2.1) |
| **TP53** | 0.064 | 0.069 | 0.6 | 1.8 (0.96-3.5) |
| **ATM** | 0.22 | 0.23 | -0.85 | 0.43 (0.11-1.7) |
| **BRAF** | 0.34 | 0.34 | 0.28 | 1.3 (0.75-2.3) |
| **DUSP2** | 0.16 | 0.17 | -0.7 | 0.5 (0.18-1.3) |
| **FAT3** | 0.16 | 0.17 | 0.42 | 1.5 (0.84-2.7) |
| **HIST1H1E** | 0.21 | 0.22 | -0.63 | 0.53 (0.2-1.4) |
| **FAM46C** | 0.15 | 0.16 | 0.35 | 1.4 (0.88-2.3) |
| **ACTG1** | 0.52 | 0.52 | -0.37 | 0.69 (0.22-2.2) |
| **NRAS** | 0.3 | 0.3 | -0.22 | 0.8 (0.53-1.2) |
| **LTB** | 0.034 | 0.039 | 0.75 | 2.1 (1-4.3) |
| **IGLL5** | 0.12 | 0.12 | 0.33 | 1.4 (0.92-2.1) |

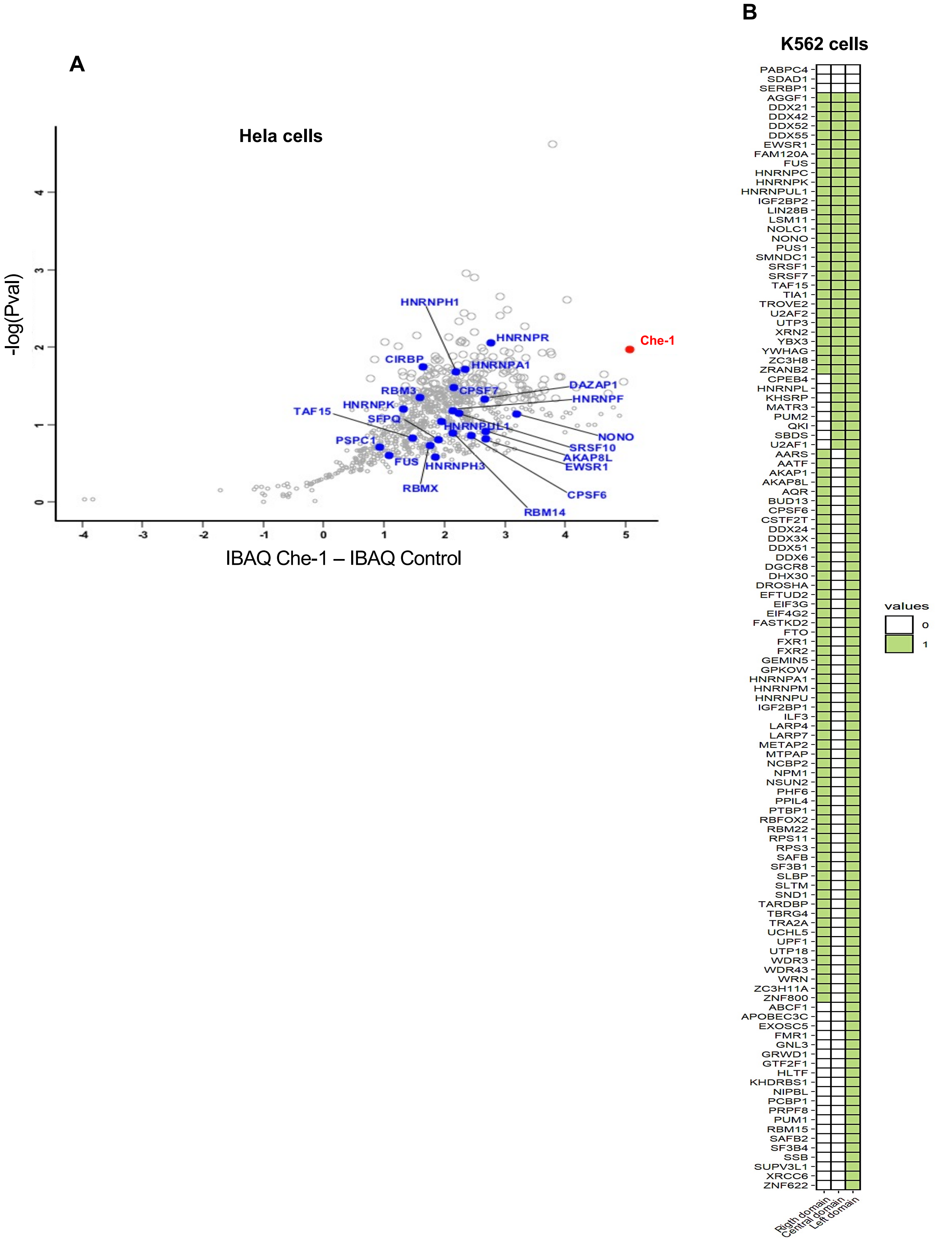

**Supplementary Figure1**

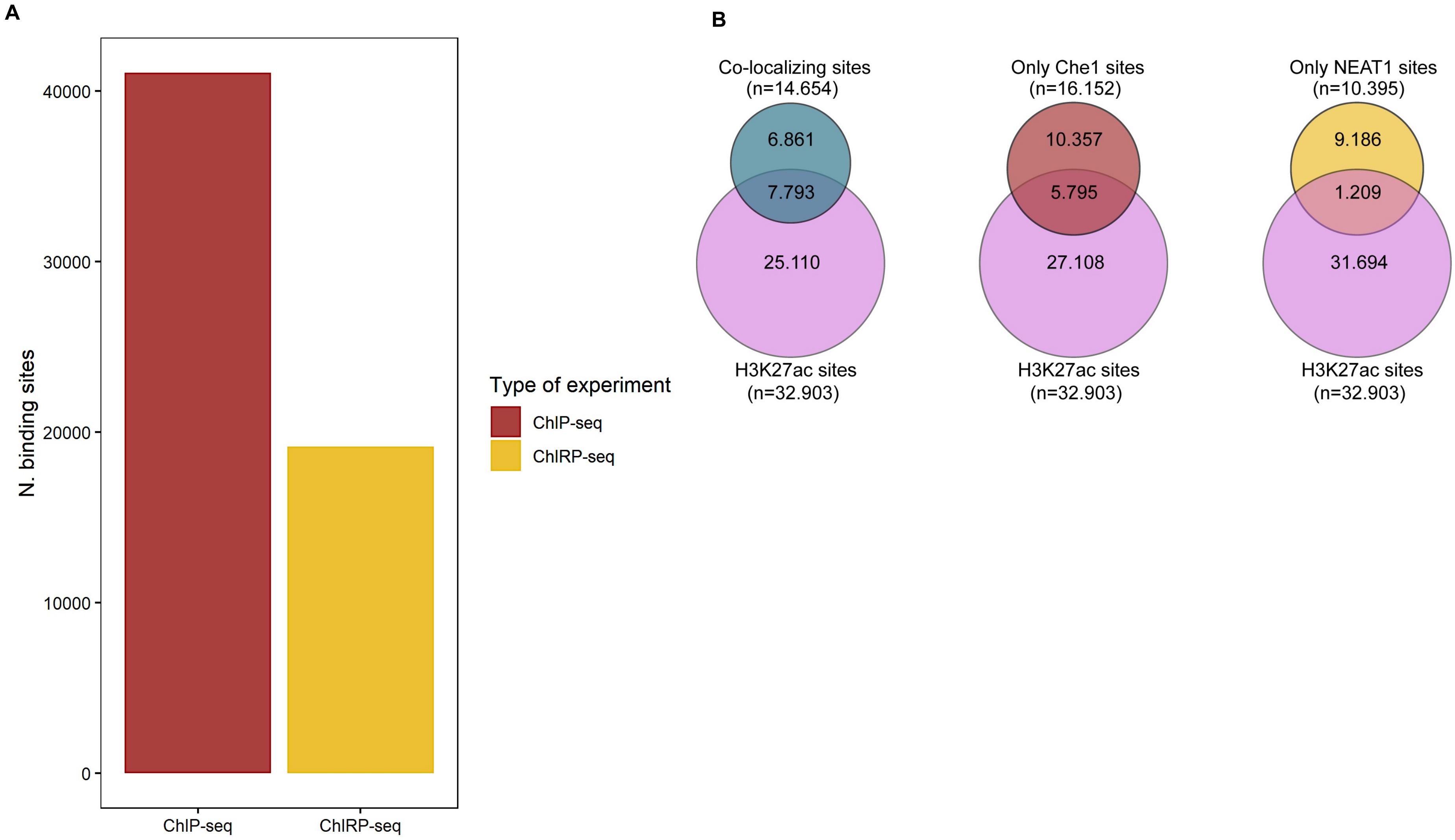

**Supplementary Figure 2**

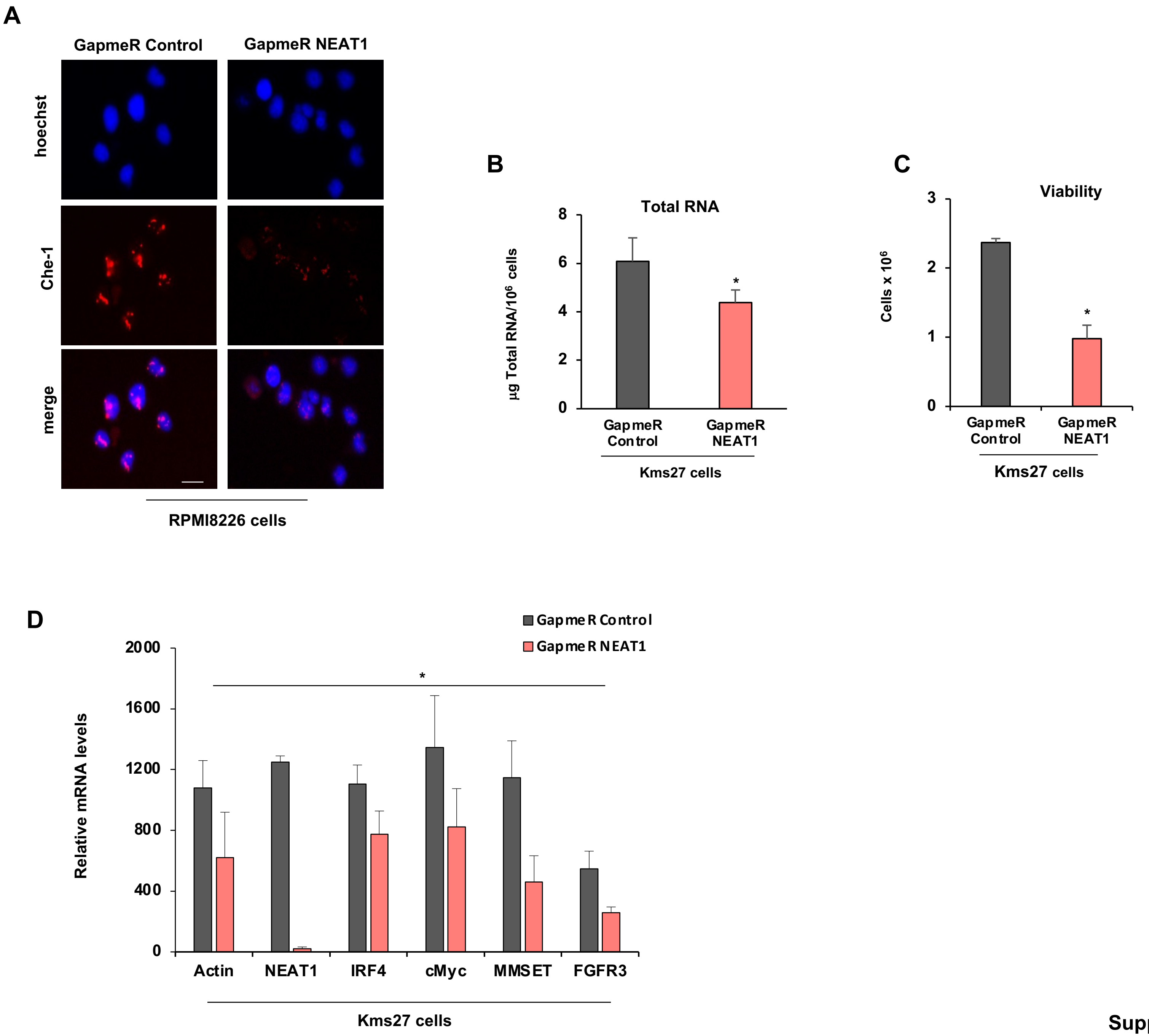

Supplementary Figure 3

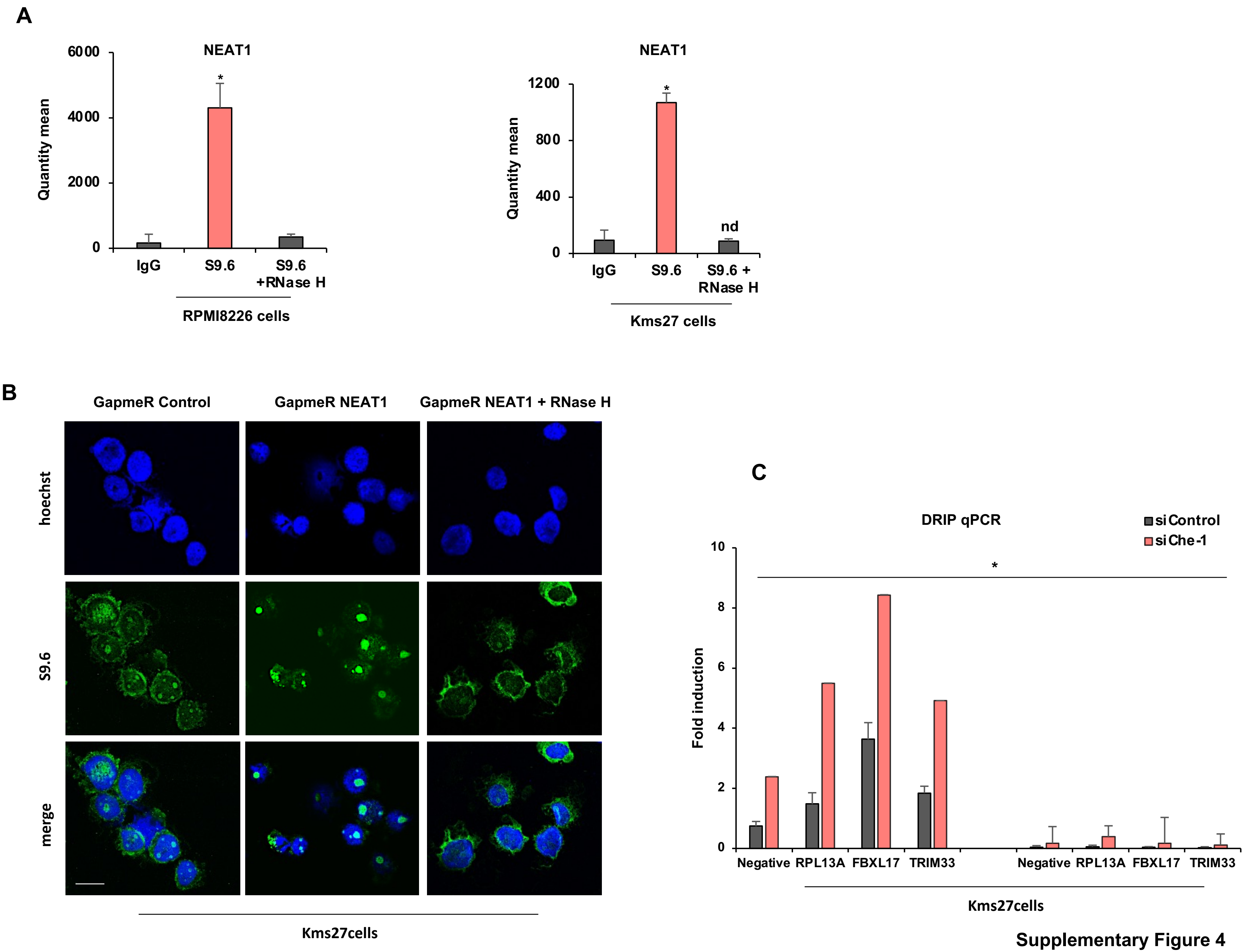

Supplementary Figure 4

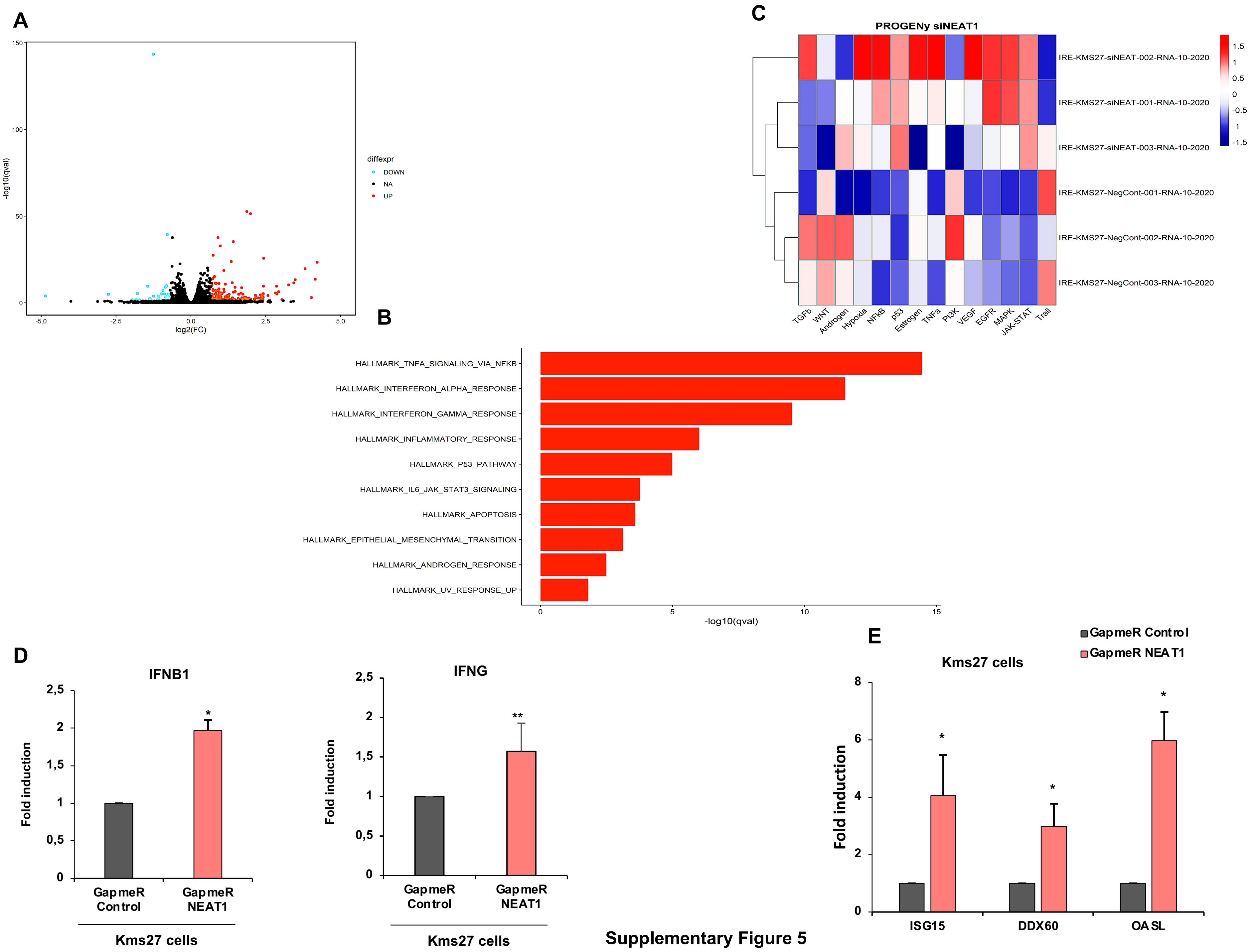

**Supplementary Figure 5**

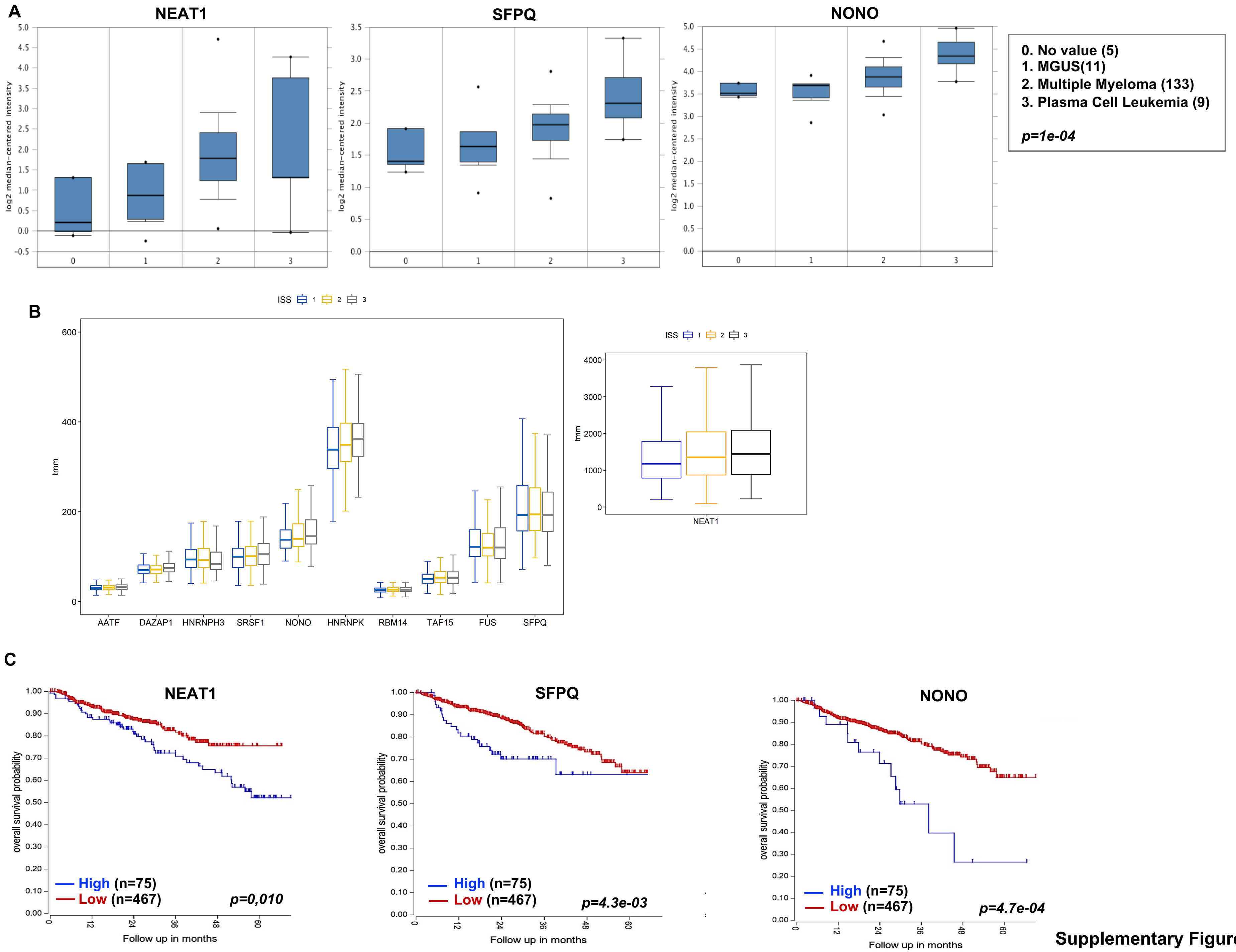

Supplementary Figure 6

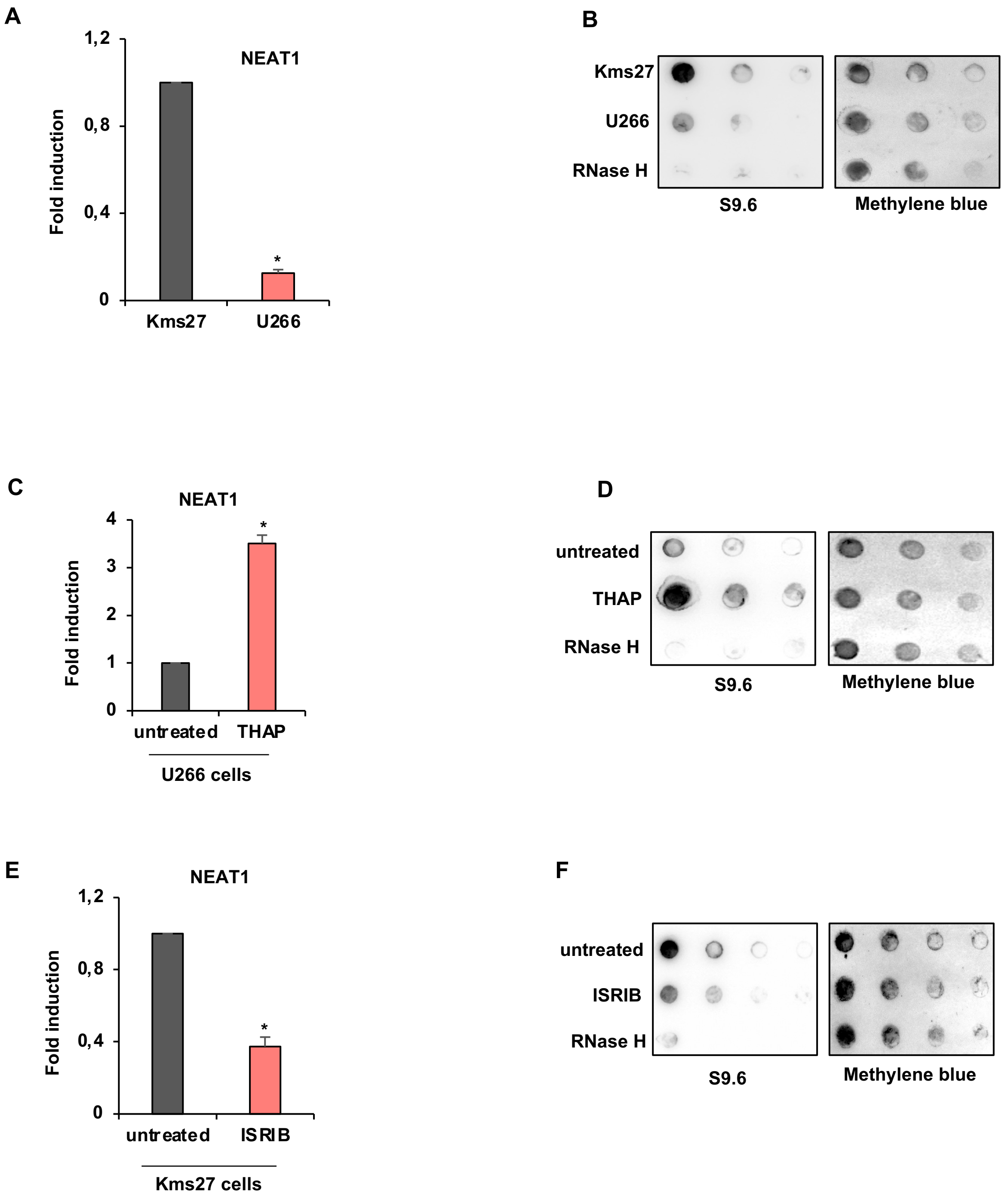

Supplementary Figure 7
